# Prevalence and drivers of nitrogen-related limitation of phytoplankton growth across space and time in Norwegian lakes

**DOI:** 10.64898/2026.05.06.723322

**Authors:** Thomas Rohrlack

## Abstract

The prevalence of nitrogen limitation and nitrogen-phosphorus co-limitation (henceforth referred to as nitrogen-related limitation) in Norwegian lakes and their relationships with atmospheric nitrogen deposition, climate, dissolved organic matter (DOM), and catchment characteristics were assessed across space and time. Routine monitoring data from 1,529 lakes in the national Vannmiljø database were analyzed for two multi-year periods (1995–2009 and 2010–2025). Limitation was inferred using the molar NO₃⁻–N/TP ratio as an indicator of dissolved inorganic nitrogen availability.

Nitrogen-related limitation was widespread in both periods and exhibited strong regional structure, with highest prevalence in northern regions and lowest prevalence in southwestern Norway. Overall prevalence increased from 31% to 38% between periods, with significant increases in western regions. Regional-scale models identified climate, forest cover, DOM, agriculture, and atmospheric nitrogen deposition as predictors of limitation probability, whereas study period *per se* and bog/peatland cover were not significant. At the local scale, atmospheric nitrogen deposition and DOM were the only consistent predictors, with substantially lower explanatory power than at the regional scale.

These results indicate that large-scale environmental gradients play a major role in shaping nutrient stoichiometry in Norwegian lakes. Because the monitoring dataset primarily represents lakes affected by human activities, the findings are particularly relevant for water management. The widespread occurrence of nitrogen-related limitation suggests that nitrogen availability may influence phytoplankton growth in many systems and that dual-nutrient management strategies addressing both nitrogen and phosphorus may be required in many regions.

## 1. Introduction

During the late 1960s and throughout the 1970s, comparative studies, time-series observations, and whole-lake fertilization experiments converged on the conclusion that phosphorus is the primary limiting nutrient for phytoplankton growth in most lakes (Edmondson, 1970; Schindler, 1974; Vollenweider, 1968). Schindler (Schindler, 1977) further argued that although nitrogen and carbon may occasionally limit primary production, biological access to large atmospheric pools of these elements ultimately drives aquatic ecosystems toward long-term phosphorus limitation of phytoplankton growth. These findings formed the basis of what later became known as the phosphorus paradigm, according to which reducing phosphorus loading represents the most effective strategy for controlling eutrophication and algal biomass in lakes.

Over the past two decades, however, growing evidence has challenged the generality of this paradigm. A global synthesis by (Elser et al., 2009) concluded that nitrogen and phosphorus limitation occur with comparable frequency across ecosystems. Experimental and observational studies across climatic regions have likewise demonstrated that nitrogen availability can strongly influence phytoplankton growth either independently or in combination with phosphorus supply (Maberly et al., 2020; Paerl et al., 2016). As a consequence, several authors have argued that effective eutrophication management may require simultaneous reductions in both nitrogen and phosphorus inputs rather than focusing exclusively on phosphorus control (Conley et al., 2009; Paerl et al., 2016).

Understanding when and where nitrogen limitation of phytoplankton growth occurs is therefore important for evaluating nutrient management strategies. Nitrogen availability can influence both the magnitude of phytoplankton production and the composition of phytoplankton communities. When nitrogen limits phytoplankton growth, reducing phosphorus inputs alone may not effectively control algal biomass. In addition, nitrogen limitation can promote the dominance of nitrogen-fixing cyanobacteria that compensate for low inorganic nitrogen concentrations by fixing atmospheric N₂ and that are often associated with harmful algal blooms and toxin production (Paerl et al., 2016). This is particularly important in phosphorus-rich lakes. Nitrogen inputs to lakes can also influence nutrient export to downstream aquatic ecosystems, where nitrogen enrichment may contribute to eutrophication in rivers, estuaries, and coastal waters (Conley et al., 2009). These considerations highlight the importance of understanding nitrogen limitation patterns when designing nutrient management strategies.

Nutrient limitation of phytoplankton growth in lakes can be assessed using experimental bioassays or inferred from nutrient stoichiometry in the water column. Although bioassays provide the most direct evidence of nutrient limitation, they are rarely available for the large spatial and temporal scales covered by routine monitoring programs. Stoichiometric indicators based on nutrient ratios are therefore widely used as screening tools for identifying potential nutrient limitation in lakes (Bergström, 2010; Ptacnik et al., 2010). Ratios based on dissolved inorganic nitrogen relative to total phosphorus (TP) have been shown to provide useful indicators of nitrogen limitation of phytoplankton growth in boreal lakes (Bergström, 2010). In contrast, ratios based on total nitrogen (TN) may be less informative because TN includes substantial pools of dissolved and particulate organic nitrogen that are often only weakly available to phytoplankton. In humic and boreal lakes, dissolved organic nitrogen can represent a large fraction of the total nitrogen pool and is closely associated with dissolved organic matter exported from catchments (Creed et al., 2018). Consequently, TN concentrations may remain high even when inorganic nitrogen available to phytoplankton is depleted during the growing season. For these reasons, dissolved inorganic nitrogen-based ratios generally provide more reliable indicators of nitrogen limitation than TN:TP in many northern lakes (Bergström, 2010; Ptacnik et al., 2010). Because ammonium measurements are not consistently available in the monitoring datasets, nitrate nitrogen (NO₃⁻–N) is often used as the most consistently measured proxy for dissolved inorganic nitrogen (de Wit et al., 2023).

In northern Europe and North America, atmospheric nitrogen deposition historically increased nitrogen availability in many lakes and shifted nutrient stoichiometry toward higher N:P ratios (Bergström et al., 2005; James J Elser et al., 2009; Hessen, 2013). More recently, atmospheric nitrogen deposition has declined substantially across Europe (Engardt et al., 2017), and long-term monitoring has documented corresponding decreases in dissolved inorganic nitrogen concentrations in lakes and streams (de Wit et al., 2023; Rogora et al., 2012). Such declines may reduce N:P stoichiometry and increase the probability of nitrogen limitation of phytoplankton growth, particularly in regions where phosphorus inputs remain elevated.

Catchment characteristics also exert strong controls on nutrient stoichiometry in lakes. Previous studies have shown that nitrogen deposition, climate, land cover, and dissolved organic matter (DOM) influence the relative availability of nitrogen and phosphorus in surface waters (de Wit et al., 2023; Hessen et al., 2009; Isles et al., 2020). Forested and peat-dominated catchments export substantial amounts of DOM, which can stimulate bacterial uptake of inorganic nitrogen, enhance denitrification, and alter internal nutrient cycling within lakes (Ask et al., 2009; Brothers et al., 2014; Weyhenmeyer and Jeppesen, 2010). Agricultural land use can further modify nutrient stoichiometry through nutrient inputs, while at the same time enhancing denitrification and other processes, which may reduce the availability of inorganic dissolved nitrogen in lakes (Finlay et al., 2013; Zhou et al., 2022). Because these drivers operate across spatial scales ranging from individual catchments to broad regional environmental gradients, identifying the dominant controls on nutrient limitation requires analyses that explicitly consider spatial scale (Burpee et al., 2022; McCullough et al., 2024; Myrstener et al., 2022; Paltsev et al., 2024).

Temporal variability adds further complexity. In many temperate and boreal lakes, dissolved inorganic nitrogen concentrations are relatively high during spring but decline during the growing season as phytoplankton and microbial communities consume available nitrogen, while TP often remains comparatively stable (Kolzau et al., 2014; Søndergaard et al., 2017). Monitoring data from Scandinavian lakes indicate that nitrogen limitation of phytoplankton growth frequently occurs during late summer when inorganic nitrogen becomes depleted (Skarbøvik et al., 2011; Søndergaard et al., 2017). Routine lake monitoring programs in northern Europe typically sample lakes intermittently during the ice-free period, most commonly between May and September, with sampling frequencies ranging from one to several observations per year. Because monitoring rarely captures the full seasonal development of nutrient limitation, pragmatic criteria are required to identify lakes that experience nitrogen depletion during this late-summer period. In the present study, nitrogen limitation was therefore inferred when the NO₃⁻–N:TP ratio fell below a threshold derived from experimental and observational studies (Bergström, 2010) during a substantial fraction of sampling occasions within a monitoring period, corresponding approximately to the most nutrient-depleted phase of the growing season.

Several environmental drivers relevant to nitrogen availability have changed markedly during recent decades. Atmospheric nitrogen deposition has declined across Europe since the 1990s (Engardt et al., 2017), while many northern lakes have experienced increasing concentrations of DOM (“browning”) associated with climate change and recovery from acidification (Meyer-Jacob et al., 2019; Weyhenmeyer et al., 2016). At the same time, some studies suggest that long-term DOM increases may be slowing or stabilizing in certain regions (Eklöf et al., 2021). Climate warming may also influence nitrogen cycling in both catchments and lakes through changes in microbial activity, mineralization rates, and hydrological transport. Together, these environmental changes suggest that nitrogen availability and nutrient limitation patterns in lakes may also be shifting over time.

The present study investigates the prevalence and environmental drivers of nitrogen as a limiting factor of phytoplankton growth in Norwegian lakes using routine monitoring data from the national Vannmiljø database. Norwegian water management is still relying on the phosphorus paradigm and a country-wide analysis of the importance of nitrogen control of phytoplankton growth in lakes with relevance for water management has not been performed. The here provided analysis focuses on the collective occurrence of nitrogen limitation and nitrogen–phosphorus co-limitation, hereafter referred to as nitrogen-related limitation, because both situations imply similar management responses involving reductions in nitrogen alongside phosphorus inputs. Furthermore, limited access to ammonium data made a separation of nitrogen limitation and nitrogen–phosphorus co-limitation unreliable (see 2.2. for more details). Using monitoring data from 1,529 lakes across two multi-year periods (1995–2009 and 2010–2025), this study addresses three questions:

1. How prevalent is nitrogen-related limitation of phytoplankton growth in Norwegian lakes, and has its prevalence changed over recent decades?
2. Which environmental gradients, including atmospheric nitrogen deposition, climate, DOM, and catchment characteristics, explain spatial variation in limitation probability?
3. Do the drivers of nitrogen-related limitation operate primarily at regional scales or at the scale of individual lakes?

To address these questions, analyses were conducted at two spatial scales. Regional analyses evaluated variation among Norwegian water regions, whereas lake-level analyses assessed drivers of limitation among individual lakes while accounting for regional clustering. Together, these approaches allow evaluation of whether nitrogen-related limitation in Norwegian lakes is predominantly structured by large-scale environmental gradients or by local catchment characteristics. It should also be noted that the monitoring data used in this study originate from the Norwegian Vannmiljø database, which primarily includes lakes that are monitored because they are affected or potentially affected by human activities. Consequently, the patterns identified here largely reflect the situation in managed or impacted systems, rather than pristine lakes.

## 2. Material and methods

### 2.1. Data collection

Lake monitoring data were obtained from the Norwegian Vannmiljø database (https://vannmiljo.miljodirektoratet.no/), which compiles water quality observations from multiple monitoring programs conducted by Norwegian environmental authorities, regional water management bodies, and research institutions. The database integrates data originating from national monitoring programs as well as regional and project-based monitoring conducted for environmental assessment and water management purposes. The dataset therefore represents routine environmental monitoring rather than measurements collected within a single standardized research program.

Data were extracted between 24 and 26 November 2025 and included surface samples or depth-integrated samples including the surface layer with measurements of NO₃⁻–N, NH₄⁺-N (if available), and TP. Only samples collected during the ice-free period were considered. Monitoring frequency varied among lakes and years because sampling was conducted within different monitoring programs and according to different monitoring objectives.

The analysis was conducted for two multi-year periods: 1995–2009 and 2010–2025. These periods were chosen to balance temporal resolution with adequate spatial coverage of lakes under irregular monitoring conditions. Many lakes were sampled only intermittently, and shorter time windows would substantially reduce the number of lakes available for analysis. The two 15-year periods therefore represent a compromise between maximizing lake coverage and allowing comparison of long-term changes in nutrient limitation patterns.

Lake color was used as primary indicator for DOM because it is routinely measured in many lakes. Where direct lake color measurements were unavailable, color was estimated from DOC or TOC measurements using regression relationships (see Section 2.3). These variables are widely used proxies for DOM concentration in freshwater systems and show strong empirical relationships across Scandinavian lakes. Lake color, DOC, and TOC data were extracted from the Vannmiljø database for the same samples as for the NO₃⁻–N, NH₄⁺-N and TP measurements.

The present study used the stoichiometric ratio of dissolved inorganic nitrogen (DIN) to TP as suggested by several authors (Bergström, 2010; Ptacnik et al., 2010). Although TN is sometimes used as a stoichiometric indicator, TN includes large pools of dissolved organic nitrogen that may not be readily available to phytoplankton, particularly in humic boreal lakes. Previous studies have therefore shown that ratios based on dissolved inorganic nitrogen are more reliable indicators of phytoplankton nitrogen limitation in such systems (Bergström, 2010).

Because the objective of this study was to identify environmental drivers of nitrogen-related limitation rather than to estimate absolute regional frequencies, representativeness of the dataset was evaluated in terms of the coverage of relevant environmental gradients rather than equal numbers of lakes in each region or period. The dataset spans wide ranges of temperature, atmospheric nitrogen deposition, land use, and DOM conditions across Norway (see Table S1), indicating that the environmental gradients relevant for nitrogen-related limitation are well represented despite differences in monitoring intensity among regions.

Atmospheric nitrogen deposition data were obtained from raster datasets produced by the European Monitoring and Evaluation Program (EMEP) representing annual deposition of oxidized and reduced nitrogen (wet and dry deposition, 10 km grid). Climate data were represented by mean annual air temperature derived from gridded datasets provided by the Norwegian Meteorological Institute (https://thredds.met.no/, 1 km grid). Spatial data describing current land use, catchments, and water management regions were downloaded from the Geonorge portal (www.geonorge.no), including the AR50 land-cover dataset, the Regine catchment database, and official Norwegian water-region boundaries (water region, see Fig. 1 for spatial distribution of water regions in Norway).

**Fig. 1:**
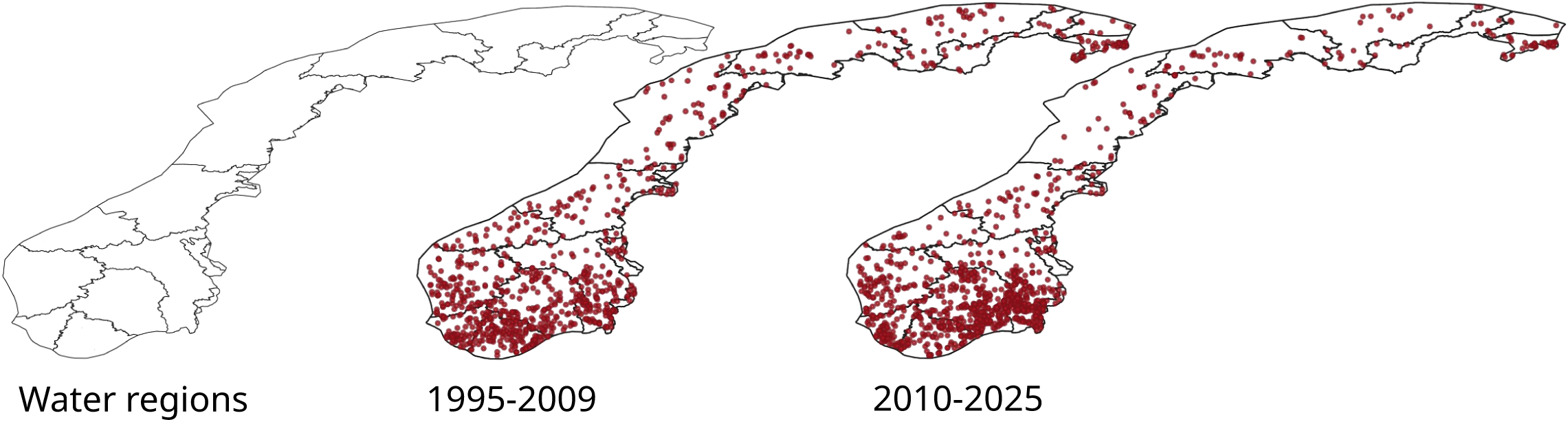
Water regions in Norway (left) and lakes included in this study (middle – first period, right second period)

Water regions are the largest spatial national management units defined under the EU Water Framework Directive. These regions represent administrative water management units rather than hydrologically independent catchments. Each water region typically includes multiple river basins and numerous lakes that may not be hydrologically connected but share similar climatic and environmental conditions. In the present study, water regions were therefore used as regional analytical units representing broad-scale environmental gradients across Norway rather than hydrologically discrete systems.

Environmental variables were extracted as spatial averages over lake catchments derived from the Regine catchment database, which provides standardized delineations of river catchments across Norway. However, individual lake catchments are not available for lakes smaller than approximately 1 km². For these lakes, the catchments of associated watercourses were used as proxies for lake catchments in order to ensure consistent spatial processing across all lakes. This approach results in catchments that may be larger than the true lake catchment and therefore represents a coarser approximation of the actual contributing area. The use of proxy catchments introduces spatial uncertainty in the representation of local land cover and environmental drivers and may increase variability in lake-level analyses. This limitation may partly explain why large-scale predictors show stronger relationships with nitrogen-related limitation than predictors evaluated at the individual-lake scale.

Monitoring data included in the Vannmiljø database originate from multiple monitoring programs conducted over several decades. As a result, sampling frequency, analytical methods, and detection limits may vary among programs and time periods, and additional datasets may have been added to the database over time. Such heterogeneity is typical for long-term environmental monitoring databases and may introduce additional variability in individual measurements. However, the large spatial and temporal coverage of the dataset reduces the influence of individual measurements or program-specific differences when evaluating broad-scale patterns. In addition, nutrient limitation was classified using threshold-based nutrient ratios rather than absolute nutrient concentrations, which reduces sensitivity to small analytical differences among laboratories or time periods. Potential variability in monitoring programs and detection limits is therefore expected primarily to introduce additional noise at the lake level rather than systematic bias in the regional patterns and environmental relationships evaluated in this study.

### 2.2. Identification of nitrogen-related limitation of phytoplankton growth

Nitrogen-related limitation was assessed using nutrient stoichiometry as a screening-level indicator. Because stoichiometric indicators infer potential nutrient limitation from nutrient ratios rather than from experimental bioassays or direct observations of ecological responses (e.g., algal growth or biomass), limitation identified in this study should be seen as inferred rather than directly measured. Stoichiometric indicators cannot demonstrate nutrient limitation of phytoplankton growth in a strict experimental sense; rather, they indicate conditions under which nitrogen availability is likely to constrain phytoplankton growth. Despite this limitation, nutrient ratios have been widely used in large-scale lake surveys where experimental assessments of nutrient limitation are not feasible (Bergström, 2010; Morris and Lewis Jr, 1988; Ptacnik et al., 2010).

Countries such as Norway, which have implemented the EU Water Framework Directive, use pressure-focused monitoring. This approach targets only those parameters that reflect the assumed dominant pressures on a given lake. Consequently, most lakes are monitored for nitrate, the primary form of nitrogen available to algae. Ammonium, by contrast, is typically included only where substantial wastewater influence or anoxic conditions are expected. This monitoring strategy had three important implications for the present study. First, the number of lakes with DIN (ammonium + nitrate) data available to the study was relatively low. During 1995–2009, DIN data were available for 261 lakes compared to 895 lakes with nitrate data; for 2010–2025, the numbers were 654 and 1,134 lakes, respectively. Second, even in lakes where ammonium was measured, sampling frequency was much lower than for nitrate (1995-2009: 3.6 individual ammonium measurements per lake, 7 for nitrate; 2009-2025: 7.5 per lake for ammonium, 12.3 per lake for nitrate). Third, available ammonium data were biased toward lakes with higher concentrations than those typically found in Norwegian lakes. The magnitude of this bias is difficult to quantify, as representative (unbiased) ammonium data from lakes relevant to water management are lacking. However, nationwide river monitoring data reported an average ammonium concentration of 14.4 µg L^-1^ in 2017 (Gundersen et al., 2019), whereas the corresponding average for lakes with ammonium data during 2010–2025 is substantially higher, at 54.5 µg L^-1^.

The limited spatial coverage, low sampling frequency, and bias toward elevated ammonium concentrations would have made analyses for the first study period unreliable at best. Ammonium data were therefore excluded, and NO₃⁻–N was used as the most consistently available indicator of dissolved inorganic nitrogen. Because ammonium was not included, the NO₃⁻–N:TP ratio represents only a subset of total dissolved inorganic nitrogen and may therefore differ from DIN:TP, particularly in systems where ammonium contributes substantially to the inorganic nitrogen pool. To assess the implications of this limitation for stoichiometric classification, a sensitivity analysis was conducted using the subset of lakes for which both ammonium and nitrate data were available. This analysis compared classifications based on molar DIN:TP and NO₃⁻–N:TP ratios using thresholds from (Bergström, 2010), where values lower or equal 3.5 (= mass ratio 1.5) indicate nitrogen limitation and values higher than 3.5 but lower or equal 7.5 (= mass ratio 3.4) indicate nitrogen–phosphorus co-limitation. As discussed in Sections 3.1. and 4, excluding ammonium affected the classification of limitation status, particularly the distinction between nitrogen limitation and nitrogen–phosphorus co-limitation. Given these constraints, nitrogen-limited and nitrogen–phosphorus co-limited lakes were not treated as separate categories. This decision reflects both the limitations of the available data and the fact that both conditions indicate that nitrogen availability may constrain phytoplankton growth and thus require management strategies addressing nitrogen inputs alongside phosphorus. Separating these categories would therefore imply a level of resolution that cannot be supported reliably by the dataset and would not substantially alter the study’s management-relevant conclusions. Instead, both categories were combined into a single group, hereafter referred to as lakes exhibiting nitrogen-related limitation of phytoplankton growth.

Following Bergström (Bergström, 2010), nitrogen-related limitation was identified when the molar NO₃⁻–N:TP ratio was <7.5. This threshold corresponds to a mass ratio of approximately 3.4 and was derived from short-term nutrient addition bioassays combined with statistical analyses of phytoplankton responses in Swedish lakes. In these experiments, DIN:TP ratios were calibrated against observed shifts between nitrogen and phosphorus limitation during the growing season, providing an empirically based basis for stoichiometric classification. Because the 7.5 threshold was originally defined for DIN:TP (NO₃^-^-N and NH₄^+^-N), its application to NO₃⁻–N:TP represents an approximation, the implications of which are evaluated using the sensitivity analysis described above. Previous studies suggest that thresholds for nitrogen-related limitation based on nutrient ratios may vary somewhat among ecosystems (Morris and Lewis Jr, 1988; Ptacnik et al., 2010), but the Bergström threshold was considered most appropriate for the present study because it was developed for lakes with climatic and geological characteristics similar to those of Norwegian lakes.

The limitation criterion was applied to individual sampling occasions. A lake was classified as exhibiting nitrogen-related limitation during a study period if the NO₃⁻–N:TP threshold was met on at least 20% of sampling occasions within that period. To ensure that classification of limitation status was based on sufficient temporal information, lakes with fewer than five sampling occasions within a study period were excluded from the analyses. This minimum requirement reduces uncertainty associated with very low sampling frequency and ensures that the 20% threshold represents a meaningful proportion of observations. Lakes not classified as exhibiting nitrogen-related limitation were not interpreted as phosphorus-limited, but rather as lakes in which nitrogen availability was not indicated to constrain phytoplankton growth based on the applied stoichiometric criterion.

The 20% threshold was chosen to capture lakes experiencing sustained seasonal nitrogen depletion rather than occasional short-term fluctuations. In Northern Europe, routine lake monitoring typically covers the ice-free period from May to September, and nitrogen limitation in boreal lakes commonly occurs during late summer when dissolved inorganic nitrogen becomes depleted. A 20% threshold therefore corresponds approximately to one month of potential nitrogen limitation within a typical monitoring season. Using this temporal threshold reduces the likelihood that lakes are classified as nitrogen-related limited due to single anomalous measurements. At the same time, lakes with only very short or episodic periods of nitrogen depletion are conservatively classified as not exhibiting nitrogen-related limitation.

Because the classification is based on the proportion of sampling occasions meeting the threshold, differences in sampling frequency among lakes do not directly bias the classification, although lakes with very few samples have inherently greater uncertainty. The use of a proportional threshold therefore allows comparison across lakes with varying monitoring intensity. Samples collected during both mixed and stratified conditions were included. Because routine monitoring typically covers the entire ice-free period, the dataset integrates seasonal variability in nutrient concentrations. The 20% threshold was specifically chosen to identify lakes that experience sustained nitrogen depletion during the growing season, rather than transient conditions associated with short-term mixing events.

### 2.3. Spatial data processing and estimation of DOM

All spatial processing and visualization were conducted using QGIS (version 3.4) and the GDAL processing toolbox. Atmospheric nitrogen deposition and climate data were obtained as gridded raster datasets and aggregated to represent average conditions for each study period. For both variables, mean values were calculated separately for each study period (1995–2009 and 2010–2025) and then intersected with catchment polygons derived from the Regine database to calculate catchment-level averages for each lake.

Land-cover information was obtained from the AR50 land-cover dataset, which provides polygon-based classifications of major land-cover types across Norway. The proportional coverage of agriculture, forest, and bog/peatland within each catchment was calculated using spatial intersection between land-cover polygons and catchment boundaries. Because AR50 represents a static land-cover map, land-cover proportions were assumed constant across both study periods. This assumption is justified by the generally low rates of land-use change in Norway, where forest and semi-natural land dominate and agricultural and built-up areas constitute a small proportion of the total land area. National statistics indicate that agricultural land extent has remained largely stable over recent decades and that major shifts in land-cover composition are limited at the spatial scale considered here (Andersson, 2025).

DOM concentrations were represented using lake color, a commonly used proxy for DOM in boreal lakes. Lake color is strongly correlated with dissolved organic carbon (DOC) across Scandinavian lakes and has therefore been widely used as an indicator of DOM concentration in regional surveys (e.g., Isles et al., 2020). For lakes lacking direct color measurements, lake color was estimated from DOC or TOC using linear regression relationships derived from lakes where both variables were available. Across both study periods, DOC, TOC, and lake color showed strong linear relationships (Figs. S1 and S2), with DOC and TOC exhibiting an approximately 1:1 relationship. Consequently, DOC and TOC were treated equivalently for the purpose of estimating lake color. This approach allowed consistent representation of DOM conditions across lakes despite differences in measured variables among monitoring programs. Mean lake color values were then calculated separately for each lake and study period using all available observations within the respective period.

### 2.4. Testing for changes in predictor variables over time

Lakes were grouped by water region. As mentioned, water regions are the largest spatial management units defined under the EU Water Framework Directive (Fig. 1). Water regions represent administrative water-management units that typically encompass several river basins and numerous lakes. In the present study, these regions were used as regional analytical units representing broad-scale environmental gradients rather than hydrologically connected systems.

For each water region, changes in key environmental predictors between the two study periods (1995–2009 and 2010–2025) were evaluated. The analysis focused on predictors expected to vary over time at regional scales, namely mean annual air temperature, atmospheric nitrogen deposition, and lake color (as a proxy for dissolved organic matter). Catchment land-cover variables were not tested for temporal change because only a single land-cover dataset was available for the study area.

Lakes were treated as independent observations within each study period, resulting in cross-sectional comparisons of environmental conditions between the two multi-year periods rather than repeated measurements of identical lakes. For each water region and predictor variable, mean values were calculated for all lakes available in each study period.

Differences in mean predictor values between the two periods were tested using Welch two-sample t-tests, which do not assume equal variances or equal sample sizes. Prior to analysis, lake color and atmospheric nitrogen deposition were log-transformed to improve distributional symmetry, whereas temperature was analyzed on the original scale.

Because multiple tests were conducted across water regions, p-values were adjusted separately for each predictor variable using the Benjamini–Hochberg false discovery rate (FDR) procedure to control for multiple comparisons.

### 2.5. Changes in prevalence of nitrogen-related limitation of phytoplankton growth over time

For each water region, changes in the prevalence of nitrogen-related limitation between the two study periods (1995–2009 and 2010–2025) were evaluated. Limitation status was treated as a binary variable, indicating whether a lake met the criterion for nitrogen-related limitation during a given study period.

Because monitoring coverage differed among lakes and years, not all lakes were sampled in both study periods. To maximize spatial coverage of environmental gradients, lakes were therefore treated as independent cross-sectional observations within each study period rather than restricting the analysis to lakes monitored in both periods. Restricting the analysis to the subset of shared lakes would substantially reduce the number of lakes available in several regions and would bias the analysis toward regions with more intensive monitoring.

For each water region, the number of lakes classified as limited and not exhibiting nitrogen-related limitation was summarized separately for each study period, resulting in 2 × 2 contingency tables (study period × limitation status). Differences in prevalence between periods were evaluated using Pearson’s χ² tests. Regions with insufficient data to construct contingency tables (e.g., absence of lakes in one period or absence of one outcome category) were excluded from these analyses.

### 2.6. Drivers of nitrogen-related limitation of phytoplankton growth: regional scale

Environmental drivers of nitrogen-related limitation were evaluated at two spatial scales: regional and lake level. The regional analysis examined how the prevalence of limitation varies among Norwegian water regions, whereas the lake-level analysis (Section 2.7) assessed how environmental predictors influence limitation probability among individual lakes.

The unit of analysis was the lake–period combination. For each lake and study period, limitation status was first determined based on the proportion of sampling occasions meeting the NO₃⁻–N:TP threshold (Section 2.4). Each lake therefore contributed at most one binary observation per study period indicating whether nitrogen-related limitation occurred during that period.

Because monitoring coverage varied among lakes and years, not all lakes were sampled in both study periods. To maximize spatial coverage across environmental gradients, lakes were therefore treated as independent cross-sectional observations within each study period rather than restricting the analysis to lakes monitored in both periods. Restricting the analysis to the subset of shared lakes would substantially reduce the number of lakes available in several water regions and bias the dataset toward areas with longer monitoring histories.

At the regional scale, lakes were grouped by water region and study period. For each region–period combination, the numbers of lakes classified as limited and not exhibiting nitrogen-related limitation were summarized. These counts represent the regional prevalence of nitrogen-related limitation. Environmental predictors (agricultural land cover, forest cover, bog/peatland cover, mean annual temperature, mean atmospheric nitrogen deposition, and mean lake color) were averaged at the region–period level.

Regional limitation probability was modeled using a binomial generalized linear model (GLM) with a logit link function, where the response variable consisted of the counts of limited versus lakes not exhibiting nitrogen-related limitation lakes in each region and study period. This formulation retains differences in regional sample size while avoiding assumptions of equal monitoring intensity across regions.

Predictor significance was evaluated using Wald tests, and explanatory power was quantified using Nagelkerke’s (Cragg–Uhler) pseudo-R². Multicollinearity among predictors was assessed using variance inflation factors (VIF). In addition, pairwise relationships among predictors were evaluated using a Spearman correlation matrix calculated across all lakes included in the dataset for each of the two study periods (Tables S2 and S3).

To assess whether relationships between environmental drivers and limitation changed over time, models including interaction terms between study period and environmental predictors were also evaluated. Because these interaction terms did not improve model fit, the final model retained only the main effects of environmental predictors.

### 2.7. Drivers of nitrogen-related limitation of phytoplankton growth: lake scale

To evaluate environmental drivers of nitrogen-related limitation at the individual-lake scale, statistical models were fitted using the limitation status of each lake–period combination as the response variable. Each lake therefore contributed at most one binary observation per study period, indicating whether nitrogen-related limitation occurred during that period.

Predictor variables were identical to those used in the regional analysis: agricultural land cover, forest cover, bog/peatland cover, mean annual air temperature, atmospheric nitrogen deposition, and mean lake color. These predictors represent catchment characteristics and large-scale environmental conditions that may influence nutrient stoichiometry and the likelihood of nitrogen-related limitation.

Because lakes located within the same water region may experience similar climatic conditions, atmospheric deposition, and land-use influences, observations from lakes within the same region cannot be assumed to be fully independent. To account for this spatial grouping, generalized estimating equations (GEE) were used. GEE models extend generalized linear models by allowing observations within predefined clusters to be correlated while still estimating population-level relationships between predictors and the response variable.

In the present analysis, water regions were treated as clustering units, meaning that lakes within the same region were allowed to be statistically correlated. An exchangeable working correlation structure was specified. This structure assumes that all lakes within the same region have approximately the same pairwise correlation in their limitation status, regardless of their geographic distance within the region. In other words, the model accounts for the possibility that lakes within a region are more similar to each other than to lakes in other regions, but it does not assume that correlation varies among individual lake pairs within a region.

The exchangeable correlation structure was chosen because it provides a simple and robust representation of within-region dependence when the detailed spatial correlation structure among lakes is unknown. Accounting for this clustering prevents underestimation of standard errors that could occur if lakes within the same region were incorrectly treated as fully independent observations.

The response variable was the binary limitation status of each lake–period combination. Predictor effects were estimated using logistic GEE models, and statistical significance was evaluated using robust Wald tests.

Because environmental conditions changed between the two study periods, GEE models were fitted separately for each period. This approach allows the influence of environmental predictors to be evaluated independently for each period without assuming that predictor effects remained constant over time.

Model performance was summarized using McFadden’s pseudo-R² and Tjur’s R², which describe overall model fit and the ability of the model to discriminate between limited and lakes not exhibiting nitrogen-related limitation. Multicollinearity among predictors was evaluated using variance inflation factors (VIF).

### 2.8. Limitations

Several limitations should be considered when interpreting the results of this study

First, nitrogen-related limitation of phytoplankton growth was identified using stoichiometric indicators rather than experimental nutrient-addition assays. Nutrient ratios provide indirect evidence of potential nutrient limitation and cannot demonstrate limitation in a strict experimental sense. Instead, they indicate conditions under which phytoplankton growth is likely to become nitrogen-limited. Such stoichiometric indicators are widely used in large-scale lake surveys because experimental assessments of nutrient limitation are not feasible for large monitoring datasets. The threshold used in this study was derived from empirical relationships between nutrient ratios and phytoplankton responses in Scandinavian lakes (Bergström, 2010), making it appropriate for the environmental conditions represented in the dataset.

Second, dissolved inorganic nitrogen was approximated using NO₃⁻–N because ammonium measurements were not consistently available in the monitoring data. This may lead to underestimation of total inorganic nitrogen availability in lakes where ammonium contributes substantially to dissolved inorganic nitrogen, and may therefore influence stoichiometric classifications of nutrient limitation. The implications of excluding ammonium were evaluated using a sensitivity analysis based on the subset of lakes with available DIN data (Sections 3.1 and 4), which allows assessment of how strongly classifications based on NO₃⁻–N:TP differ from those based on DIN:TP.

Third, catchment characteristics were derived from the Regine catchment database, which does not provide individual lake catchments for lakes smaller than approximately 1 km². For these lakes, larger proxy catchments associated with nearby watercourses were used. This approach may reduce the accuracy of catchment land-cover estimates for some lakes and may partly contribute to weaker relationships between local land-cover predictors and limitation status. However, this uncertainty is unlikely to affect the broad spatial gradients analyzed at regional scales.

Finally, the monitoring data compiled in the Vannmiljø database originate from multiple monitoring programs conducted over several decades. Sampling frequency, analytical methods, and detection limits may therefore vary among lakes and time periods. Such heterogeneity is typical of large environmental monitoring databases and may introduce additional variability in individual measurements.

Because the analyses focus on broad spatial gradients across a large number of lakes, these sources of variability are expected primarily to introduce additional noise rather than systematic bias in the observed relationships. The consistency of spatial patterns across both regional and lake-level analyses further suggests that the main conclusions are robust to these methodological uncertainties.

## 3. Results

### 3.1. Sensitivity of stoichiometric classification to exclusion of ammonium

To evaluate the effect of excluding ammonium, classifications based on NO₃⁻–N:TP were compared with classifications based on DIN:TP for the subset of lakes with available ammonium data. In the period 1995–2009, DIN:TP classified 6.9% of lakes as nitrogen-limited and 16.9% as nitrogen–phosphorus co-limited, whereas NO₃⁻–N:TP classified 23.3% and 9.9% of lakes into these categories, respectively. In 2010–2025, corresponding values were 11.9% and 14.7% for DIN:TP, and 29.8% and 6.2% for NO₃⁻–N:TP. When nitrogen limitation and nitrogen–phosphorus co-limitation were combined into a single category of nitrogen-related limitation, agreement between classifications based on DIN:TP and NO₃⁻–N:TP was high (89.8% for 1995–2009 and 90.0% for 2010–2025). Differences between the two approaches occurred primarily in the distinction between nitrogen limitation and nitrogen–phosphorus co-limitation, whereas classification of lakes as exhibiting nitrogen-related limitation versus phosphorus limitation was largely consistent.

### 3.2. Dataset characteristics (for all NO₃⁻–N:TP based analysis)

The following section summarizes the characteristics of the dataset used for the main analyses, based on NO₃⁻–N and TP measurements. The compiled dataset used for all subsequent analyses (i.e., based on NO₃⁻–N and TP measurements) included 1,529 lakes distributed across 13 of the 16 Norwegian water regions. The spatial distribution is shown in Fig. 1. Across both study periods, a total of 23,503 individual water samples containing paired NO₃⁻–N and TP measurements were available for analysis. More than 70% of all samples in both periods were collected between May and September, corresponding to the main growing season. In the first study period (1995–2009), 895 lakes met the data requirements for analysis, while 1,134 lakes were available in the second period (2010–2025). Approximately 450 lakes were sampled in both study periods, whereas the remaining lakes were present in only one period due to changes in monitoring coverage over time.

Monitoring intensity varied among lakes because data were compiled from multiple monitoring programs. On average, lakes had observations in 2.2 distinct years during the first period and 3.1 distinct years during the second period, indicating slightly increased monitoring coverage in the later period.

The dataset spans wide environmental gradients in temperature, atmospheric nitrogen deposition, catchment land cover, and DOM conditions (Table S1). Relationships among the DOM indicators used in this study were strong: DOC, TOC, and lake color showed highly significant linear relationships (Figs. S1 and S2). DOC and TOC were approximately proportional across lakes (Fig. S1), supporting their use as interchangeable proxies for estimating lake color where direct measurements were unavailable.

### 3.3. Changes in predictor variables over time

Environmental predictors exhibited substantial variation across water regions and between the two study periods (Figs. 2-4). Mean annual air temperature increased between the periods 1995–2009 and 2010–2025 in many water regions (Figs. 2a and 3). Regional comparisons indicated significant increases in several regions, consistent with the overall warming trend observed across Norway (Figs. 2a and 3).

**Fig. 2:**
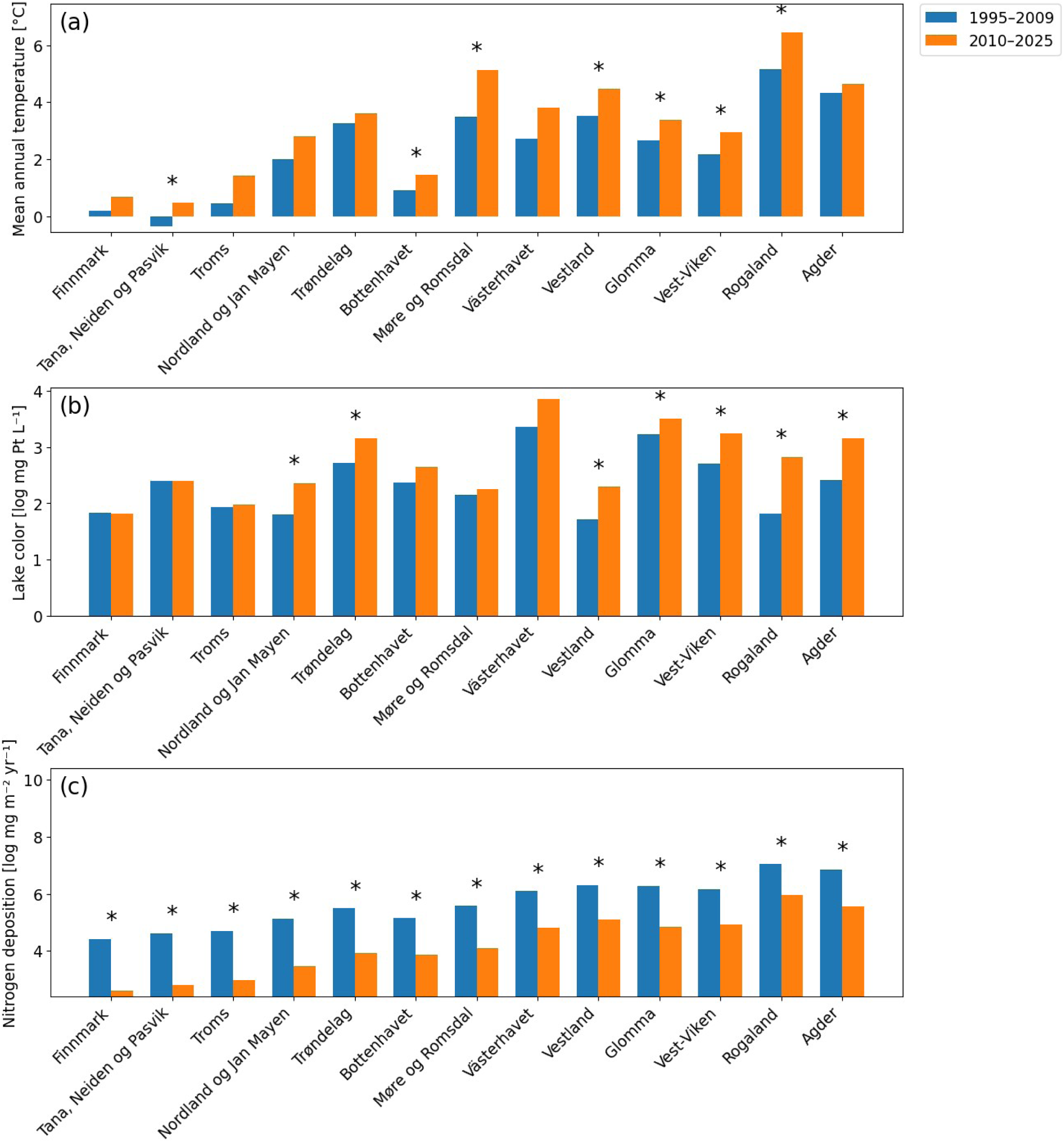
Mean annual temperature, mean lake color, and mean atmospheric nitrogen deposition in water regions during the first (1995-2009) and second study period (2010-2025). Asterisks indicate significant changes from the first to the second study period. Water regions are ordered from north (left) to south (right) and span a latitudinal gradient from approximately 58°N to 71°N.

**Fig. 3:**
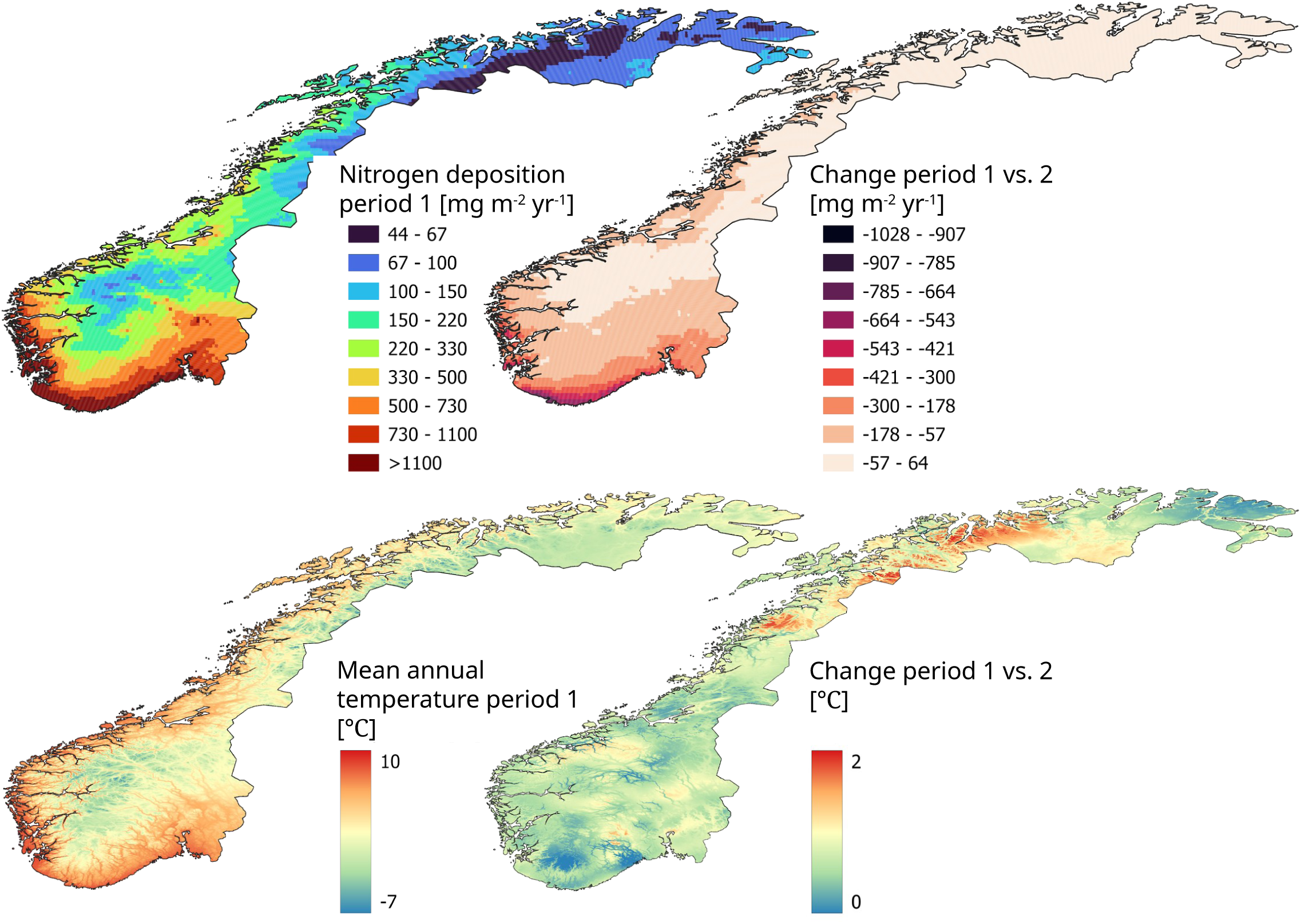
Mean atmospheric nitrogen deposition and mean annual temperature during the first study period (left, 1995-2009) and change from the first to the second study period (right, 2010-2025).

**Fig. 4:**
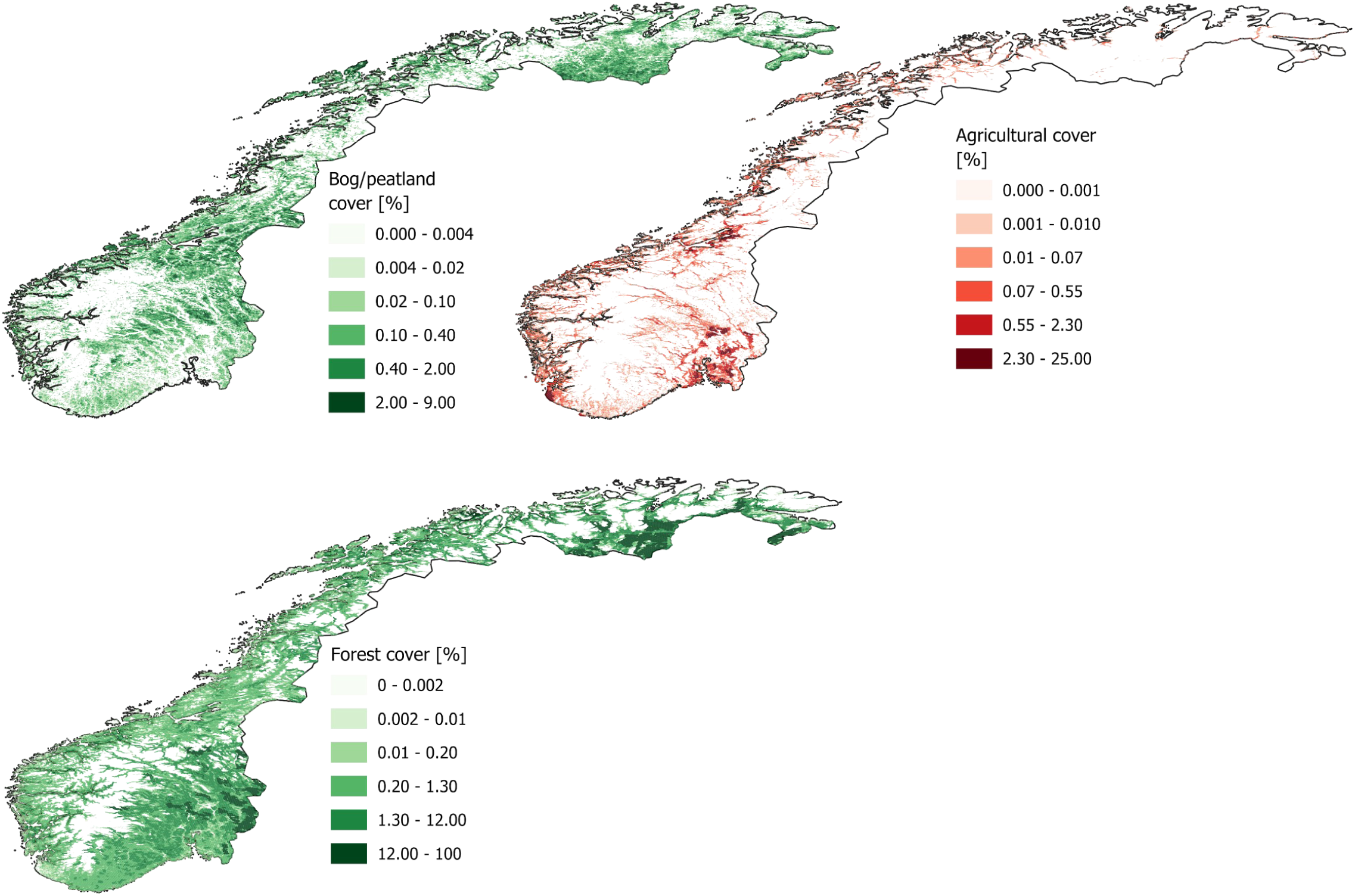
Spatial distribution of bog/peatland cover, forest cover and agricultural areas in Norway.

Changes in lake color, used here as a proxy for DOM, were more heterogeneous (Fig. 2b). Some water regions showed increases in color between the two periods, whereas others showed relatively little change.

In contrast, atmospheric nitrogen deposition generally decreased between the two study periods (Figs. 2c and 3). This decline was observed across all water regions, although the magnitude of change varied regionally. The observed decrease in atmospheric nitrogen deposition is consistent with previously documented declines in nitrate concentrations in Norwegian surface waters (de Wit et al., 2023).

Statistical comparisons of predictor values between periods were conducted using region-specific Welch two-sample t-tests with false discovery rate correction. The resulting statistical data are summarized in Table S4.

### 3.4. Prevalence of nitrogen-related limitation of phytoplankton growth

In the first study period (1995–2009), 31 % of lakes were classified as exhibiting nitrogen-related limitation, whereas 38% of lakes met the criterion in the second period (2010–2025). These values indicate an increase in the overall prevalence of nitrogen-related limitation between the two periods (Fig. 5).

**Fig. 5:**
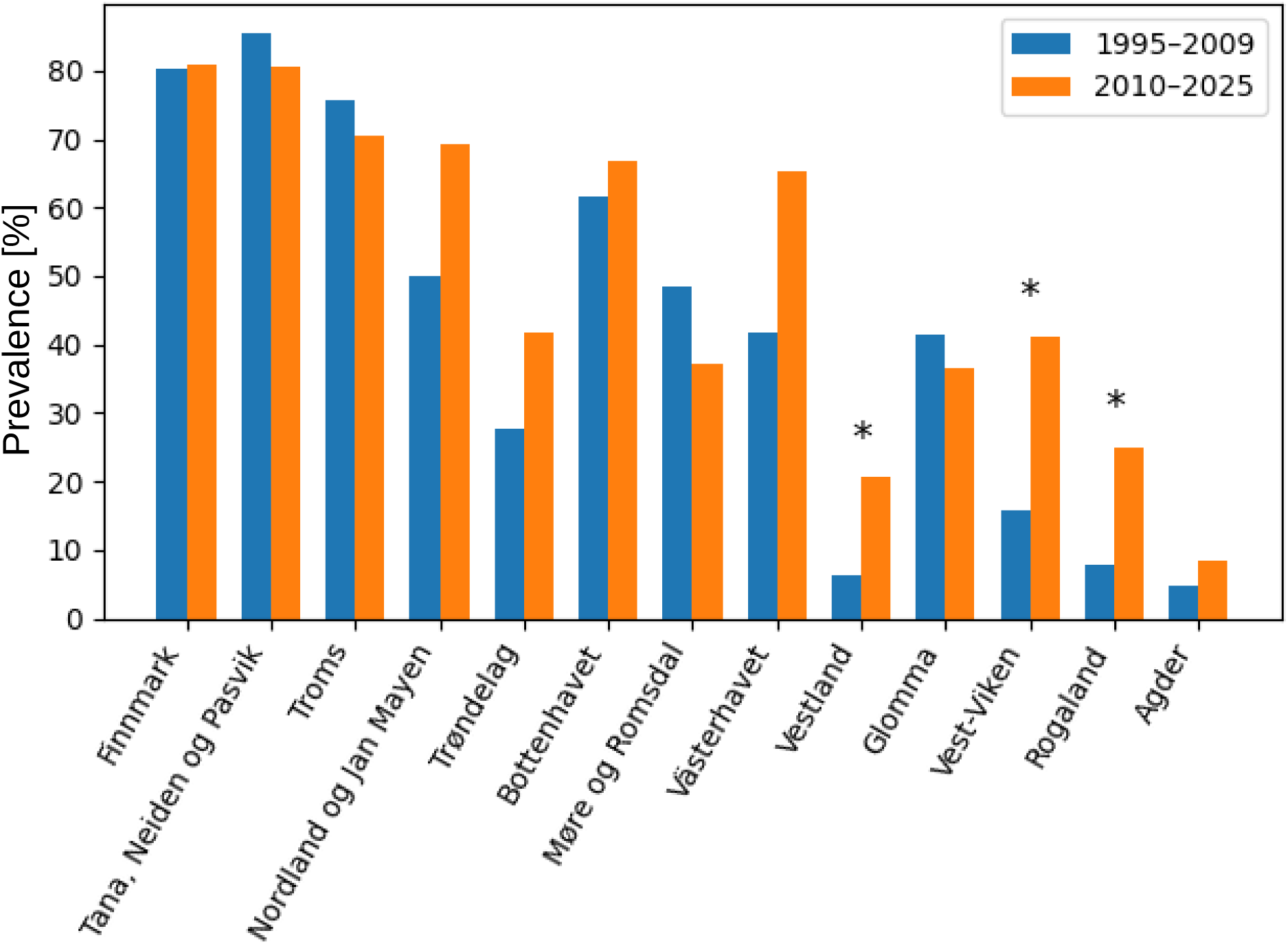
Prevalence of nitrogen-related limitation of phytoplankton growth in water regions during study periods one (1995-2009) and two (2010-2025). Asterisks indicate significant changes from the first to the second study period. Water regions are ordered from north (left) to south (right) and span a latitudinal gradient from approximately 58°N to 71°N.

The prevalence of nitrogen-related limitation varied substantially among water regions, exhibiting a north-south gradient (Fig. 5). Some regions showed consistently high proportions of limited lakes in both periods, whereas others exhibited relatively low prevalence. In three western regions (Vestland, Vest-Viken, Rogaland) the proportion of lakes classified as limited increased significantly between the two study periods, while other regions showed little change (Fig. 5).

Regional differences in limitation prevalence were statistically evaluated using contingency tests and logistic regression models. The resulting regional prevalence estimates and statistical comparisons are summarized in Table S5.

### 3.5. Regional-scale drivers of nitrogen-related limitation of phytoplankton growth

Regional variation in nitrogen-related limitation was strongly associated with several environmental predictors (Table 1). The binomial GLM explained a substantial proportion of the variation in regional limitation prevalence (Nagelkerke pseudo-R² ≈ 1).

**Table 1:** Results of the GLM analysis to identify drivers of nitrogen-related limitation of phytoplankton growth at the regional scale.

| Predictor | Estimate | Standard error | p-value | VIF |
| --- | --- | --- | --- | --- |
| Intercept | 1.670 | 0.238 | 0.000 | - |
| Period (1995-2009 vs 2010-2025) | 0.069 | 0.239 | 0.774 | 2.83 |
| Mean annual temperature | -0.572 | 0.066 | 0.000 | 3.21 |
| Atmospheric N deposition | -0.002 | 0.001 | 0.003 | 4.51 |
| Agriculture | 0.059 | 0.017 | 0.000 | 2.18 |
| Forest | -0.028 | 0.005 | 0.000 | 2.13 |
| Bog / peatland | 0.008 | 0.024 | 0.745 | 1.41 |
| Lake color (proxy for DOM) | 0.029 | 0.007 | 0.000 | 1.95 |

Among the predictors considered, lake color and relative importance of agriculture in the catchment showed positive relationships with nitrogen-related limitation (Table 1). In contrast, mean annual temperature, atmospheric nitrogen deposition, and forest cover were negatively linked to nitrogen-related limitation.

Multicollinearity among predictors was low (all VIF < 5, Table 1), and pairwise correlations among predictors were moderate (Tables S2 and S3), indicating that the effects of individual predictors could be evaluated without strong confounding. However, mean annual temperature and atmospheric nitrogen deposition showed VIF values and spearman correlations that indicate a degree of covariance that need to be considered when interpreting results (Tables 1, S2 and S3, see discussion for more detail).

Models including interaction terms between study period and environmental predictors did not show significant interaction effects, indicating that the relationships between predictors and nitrogen-related limitation were broadly consistent between the two study periods (Table 1). Consequently, the final regional model retained only the main effects of environmental predictors.

### 3.6. Lake-scale drivers of nitrogen-related limitation of phytoplankton growth

Model coefficients and statistical tests for all predictors are summarized in Tables 2 and 3. Environmental predictors also showed significant associations with nitrogen-related limitation at the lake scale (Tables 2 and 3). The logistic GEE models explained a moderate proportion of the variation in limitation status among lakes. For 1995–2009, the local-scale model showed good discriminatory power (McFadden’s pseudo-R² = 0.23; Tjur’s R² = 0.24). In 2010–2025, explanatory power was lower (McFadden’s pseudo-R² = 0.10; Tjur’s R² = 0.13), indicating greater unexplained heterogeneity among lakes in the later period.

**Table 2:** Results of the GEE analysis to identify drivers of nitrogen-related limitation of phytoplankton growth at the lake scale for the study period 1995-2009.

| Predictor | Estimate | Standard error | p-value | VIF |
| --- | --- | --- | --- | --- |
| Intercept | -0.336 | 0.341 | 0.324 | - |
| Mean annual temperature | -0.017 | 0.053 | 0.743 | 3.05 |
| Atmospheric N deposition | -0.002 | 0.001 | 0.000 | 4.75 |
| Agriculture | 0.034 | 0.027 | 0.171 | 1.51 |
| Forest | -0.013 | 0.006 | 0.051 | 1.67 |
| Bog / peatland | 0.081 | 0.047 | 0.084 | 1.40 |
| Lake color (proxy for DOM) | 0.035 | 0.008 | 0.000 | 1.21 |

**Table 3:**
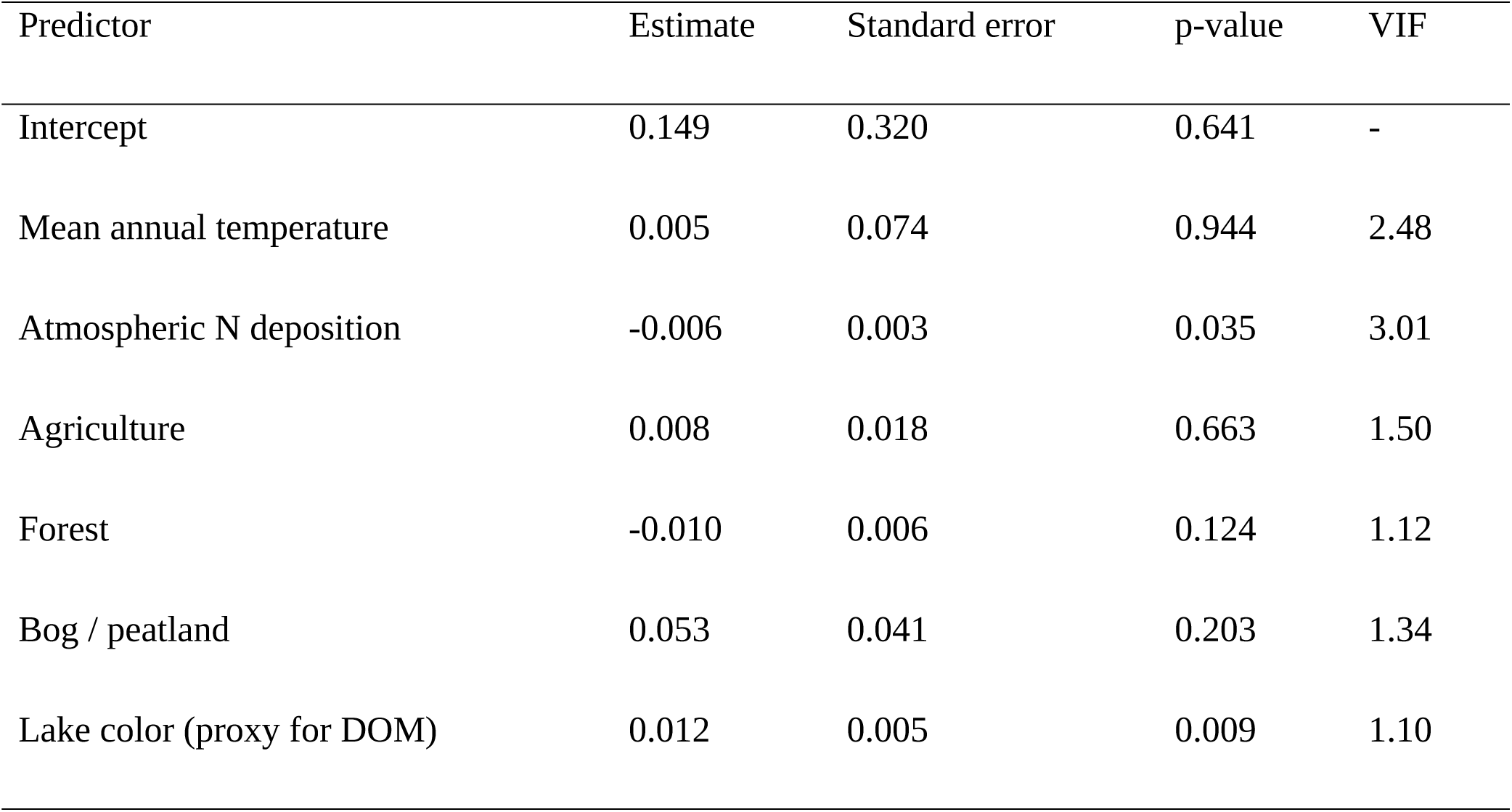
Results of the GEE analysis to identify drivers of nitrogen-related limitation of phytoplankton growth at the lake scale for the study period 2010-2025.

| Predictor | Estimate | Standard error | p-value | VIF |
| --- | --- | --- | --- | --- |
| Intercept | 0.149 | 0.320 | 0.641 | - |
| Mean annual temperature | 0.005 | 0.074 | 0.944 | 2.48 |
| Atmospheric N deposition | -0.006 | 0.003 | 0.035 | 3.01 |
| Agriculture | 0.008 | 0.018 | 0.663 | 1.50 |
| Forest | -0.010 | 0.006 | 0.124 | 1.12 |
| Bog / peatland | 0.053 | 0.041 | 0.203 | 1.34 |
| Lake color (proxy for DOM) | 0.012 | 0.005 | 0.009 | 1.10 |

Consistent with the regional analysis, lake color was positively associated with the probability of nitrogen-related limitation, indicating that lakes with higher DOM concentrations were more likely to exhibit limitation (Tables 2 and 3).

Also consistent with the regional analysis, atmospheric nitrogen deposition was negatively associated with limitation probability, suggesting that lakes located in regions receiving higher nitrogen inputs were less likely to exhibit nitrogen-related limitation (Tables 2 and 3).

## 4. Discussion

A key methodological consideration in this study is the use of NO₃⁻–N as a proxy for dissolved inorganic nitrogen due to limited availability of ammonium data. The sensitivity analysis provides important context for interpreting the resulting stoichiometric classifications. While NO₃⁻–N:TP yielded higher proportions of lakes classified as nitrogen-limited and lower proportions classified as nitrogen–phosphorus co-limited compared with DIN:TP, this pattern indicates that excluding ammonium tends to shift classifications from nitrogen–phosphorus co-limitation toward nitrogen limitation. Importantly, overall agreement between the two approaches remained high (approximately 90%) when both categories were combined into a single class of nitrogen-related limitation. This indicates that the exclusion of ammonium primarily affects the distinction between nitrogen limitation and nitrogen–phosphorus co-limitation rather than the identification of systems in which nitrogen availability may constrain phytoplankton growth.

Given this pattern, separating nitrogen limitation from nitrogen–phosphorus co-limitation would imply a level of resolution that is not robustly supported by the available monitoring data. Moreover, both conditions have similar implications for nutrient management, as they indicate that nitrogen availability can influence phytoplankton growth alongside phosphorus. The combined category of nitrogen-related limitation used here therefore provides a more robust and management-relevant representation of nutrient limitation patterns at the scale of this study. These results support the use of NO₃⁻–N:TP as a pragmatic proxy for assessing nitrogen-related limitation in large-scale monitoring datasets where full DIN measurements are unavailable.

This study demonstrates that nitrogen-related limitation of phytoplankton growth is widespread in Norwegian lakes and exhibits strong spatial organization. Across both study periods, nitrogen-related limitation was most prevalent in northern water regions and least prevalent in southwestern Norway, indicating that large-scale gradients dominate over local variability. The substantially higher explanatory power of regional-scale models compared with lake-level models further supports the conclusion that nitrogen-related limitation is primarily structured by broad-scale drivers rather than by fine-scale catchment characteristics alone. Importantly, regional models showed no interaction between study period and environmental predictors, indicating that temporal changes in limitation prevalence were associated with shifts in predictor values rather than with changes in the underlying relationships between predictors and limitation probability.

Norway represents an informative case study for evaluating the long-term validity of the phosphorus paradigm. In contrast to a growing body of literature emphasizing the importance of dual nutrient control (Conley et al., 2009; James J Elser et al., 2009; Paerl et al., 2016), Norwegian lake management still largely focuses on phosphorus reduction. The present analysis suggests that this focus may be insufficient in many systems. Rather than indicating the expected general shift toward stronger phosphorus limitation, the results show that nitrogen-related limitation of phytoplankton growth has remained common and in some water regions has increased over the last three decades. This pattern is consistent with the view that water management practices in Norway and large-scale environmental drivers of nutrient availability may act in opposing directions, partially offsetting one another.

At the same time, the present study identifies large-scale patterns in nutrient stoichiometry rather than directly testing management outcomes. The observed patterns reflect both natural environmental gradients and human influence on nutrient inputs. Consequently, the results support the relevance of nitrogen for lake management, but they do not by themselves demonstrate the effectiveness of specific management measures in individual lakes.

Climate, represented by mean annual temperature, emerged as a strong predictor of nitrogen-related limitation at the regional scale, but with a negative relationship. Regions with higher mean annual temperature showed a lower probability of nitrogen-related limitation. This result does not contradict earlier work identifying atmospheric nitrogen deposition as a major driver of lake N:P stoichiometry (Bergström et al., 2005; Crowley et al., 2012; James J. Elser et al., 2009, 2009). In Norway, strong north–south gradients in temperature and nitrogen deposition covary spatially, making their effects difficult to disentangle statistically. Warmer climates may enhance nitrogen mobilization in catchments (Bai et al., 2013; Rustad et al., 2001) and nitrogen recycling within lakes through increased mineralization and microbial activity (Brauer et al., 2013; Kellerman et al., 2014; Liikanen et al., 2002; Porcal et al., 2015), thereby reducing the probability of inorganic nitrogen depletion during the growing season. Atmospheric nitrogen deposition nevertheless exerted a consistent influence across spatial scales, with higher deposition reducing the probability of nitrogen-related limitation in both regional and lake-scale analyses. The decline in deposition observed across all water regions between the two study periods is therefore a plausible driver of the increasing prevalence of nitrogen-related limitation, particularly in regions where deposition had previously buffered lakes against nitrogen scarcity.

Forest cover was associated with a reduced probability of nitrogen-related limitation at the regional scale. While this appears to contrast with earlier Norwegian studies reporting lower NO₃⁻-N/TP ratios in forested catchments (de Wit et al., 2023; Hessen, 2013; Hessen et al., 2009), those studies were largely based on autumn sampling. In autumn, low water temperatures, reduced residence time, and low exposure to high light conditions limit in-lake mineralization of dissolved organic nitrogen derived from forest soils (Porcal et al., 2015). Autumn measurements therefore mainly represent the impact of catchment process and hydrology. In the present study, most observations were collected during the growing season, when higher temperatures and light availability may promote in-lake mineralization of dissolved organic nitrogen (Porcal et al., 2015). Under such conditions, forest-derived organic matter may increase inorganic nitrogen availability within lakes and thereby reduce the probability of nitrogen-related limitation of phytoplankton growth. The present results therefore do not necessarily contradict earlier findings, but instead emphasize the importance of seasonality and DOM when assessing nutrient limitation.

Agricultural land use increased the probability of nitrogen-related limitation at the regional scale, consistent with global evidence that cultural eutrophication often shifts lakes toward nitrogen limitation or nitrogen–phosphorus co-limitation (Finlay et al., 2013; Scott et al., 2019; Yan et al., 2016; Zhou et al., 2022). This shift can be explained by a combination of factors, including relatively low N:P stoichiometry of agricultural runoff, enhanced nitrogen losses via denitrification, increased internal phosphorus loading, and biological N₂ fixation that is insufficient to compensate for declining N:P ratios. Compared with other European countries, Norway exhibits a relatively high nitrogen loss from agricultural areas, while phosphorus loss exceeds the EU average by a wide margin (see https://ec.europa.eu/eurostat/databrowser/, (Hellsten et al., 2019)). Together, these patterns suggest a particularly low N:P stoichiometry of agricultural runoff in Norway, which may further elevate the risk of nitrogen-related limitation in agricultural regions.

Lake color, used here as a proxy for DOM, emerged as a robust predictor across spatial scales and time periods. Higher color was consistently associated with a higher probability of nitrogen-related limitation, in agreement with previous studies (Isles et al., 2020). This relationship is commonly attributed to enhanced bacterial uptake of inorganic nitrogen, increased denitrification, and stimulation of internal phosphorus loading in humic systems, all of which act to lower inorganic N:P ratios (Ask et al., 2009; Brothers et al., 2014; Weyhenmeyer and Jeppesen, 2010). At the same time, lake color is not a perfect measure of DOM concentration because iron can strongly influence water color independently of dissolved organic carbon (Kritzberg and Ekström, 2012). Lake color should therefore be interpreted here as an integrated indicator of catchment-derived brown-water conditions rather than as a direct measure of DOM alone. This limitation does not weaken the observed association but suggests caution in assigning the mechanism exclusively to DOM quantity.

The present analysis is based on nutrient ratios and therefore does not explicitly separate whether changes in nitrogen-related limitation are driven primarily by changes in nitrate, total phosphorus, or both. However, previous studies from Norway and other parts of northern Europe indicate that declining atmospheric nitrogen deposition has led to substantial reductions in nitrate concentrations in surface waters (de Wit et al., 2023), while trends in total phosphorus are more variable and may reflect a combination of reduced inputs, climate-driven changes, and increasing dissolved organic matter (“browning”). The increase in nitrogen-related limitation observed in this study is therefore consistent with a general decline in inorganic nitrogen availability relative to phosphorus, although the relative contribution of changes in NO₃⁻–N and TP likely varies among regions.

Because the focus of this study is on large-scale patterns in nutrient stoichiometry rather than on absolute nutrient concentrations, the analysis emphasizes ratios rather than individual nutrient trends. Nevertheless, absolute concentrations remain important for management, and future analyses combining stoichiometric indicators with explicit trends in NO₃⁻–N and TP would provide additional insight into the mechanisms underlying changes in nutrient limitation.

Precipitation may also influence nutrient concentrations through dilution effects and catchment export processes, but was not included due to lack of consistent spatial datasets covering both study periods. Including hydrological variables represents an important direction for future work.

The contrast between strong regional-scale relationships and weaker lake-level relationships highlights the scale dependence of nitrogen-related limitation. At the lake scale, only atmospheric nitrogen deposition and lake color consistently influenced limitation probability, and model explanatory power was moderate. This reduced performance does not necessarily imply that local processes are unimportant. Rather, local analyses must explain substantially more variability associated with lake morphology, hydrology, internal nutrient cycling, and measurement uncertainty, including uncertainty introduced by proxy catchments and irregular monitoring. Regional analyses, by contrast, average over much of this local heterogeneity and therefore more clearly reveal broad-scale environmental gradients. The consistency of the main predictor effects across both spatial scales nevertheless supports the conclusion that these gradients play an important role in shaping nitrogen-related limitation across Norwegian lakes. Although land-cover variables were treated as static across the two study periods, this is unlikely to bias the results because land-use change in Norway is generally slow and small relative to the large-scale environmental gradients examined here.

From a management perspective, the results support explicit consideration of nitrogen together with phosphorus, particularly in catchments influenced by agriculture and in regions where declining atmospheric nitrogen deposition and increasing brownification favor lower inorganic N:P ratios. However, the present results do not imply that a uniform country-wide management prescription should replace local assessment. Rather, they suggest that the risk of nitrogen-related limitation is sufficiently widespread that nitrogen should no longer be ignored in Norwegian lake management. In practice, this means that dual-nutrient management is likely to be most relevant in nutrient-enriched and agriculturally influenced systems, while regionally structured large-scale drivers should also be considered when prioritizing management actions.

Finally, it is important to emphasize that the present study is based on nitrogen-related limitation of phytoplankton growth derived from stoichiometric indicators rather than direct experimental evidence. Such indicators are well established for screening-level assessments, but they cannot capture short-term ecological dynamics, species-specific responses, or bloom formation directly. Future work combining large-scale monitoring with targeted bioassays and phytoplankton response data would therefore help to further refine the ecological interpretation of inferred nutrient limitation and strengthen the basis for management decisions.

## 5. Conclusions

This study provides the first nationwide assessment of nitrogen-related limitation of phytoplankton growth in Norwegian lakes that are significantly affected or potentially significantly affected by human activities in the catchment. The results show that nitrogen-related limitation is widespread and that its spatial distribution is strongly associated with large-scale environmental gradients, particularly lake color, atmospheric nitrogen deposition, and climate.

The consistency of these patterns across both regional and lake-scale analyses suggests that broad environmental gradients play a key role in shaping nutrient stoichiometry in Norwegian lakes. At the same time, substantial variability among individual lakes highlights the influence of local ecological processes and the complexity of nutrient dynamics at smaller spatial scales.

Because the monitoring data originate primarily from lakes affected by human activities, the results are particularly relevant for water management and eutrophication control. The widespread occurrence of nitrogen-related limitation indicates that nitrogen availability may influence phytoplankton growth in many systems and that dual-nutrient management strategies, addressing both nitrogen and phosphorus inputs, may be appropriate in some regions. However, the strong regional variability observed in this study suggests that nutrient management strategies should be adapted to regional environmental conditions rather than applied uniformly at the national scale.

Future work combining large-scale monitoring data with experimental assessments of nutrient limitation and detailed analyses of nutrient management histories would help further clarify the role of nitrogen in controlling phytoplankton dynamics and eutrophication in Norwegian lakes.

## 6. Acknowledgments

I am gratefully acknowledge the many anonymous individuals and institutions who have contributed monitoring data to the Norwegian Vannmiljø database. Their long-term efforts in data collection, quality control, and data management made this study possible. I also acknowledge the Norwegian Environment Agency and associated authorities for maintaining and providing public access to the database.

## 7. Author contributions

TR: Conceptualization, formal analysis, investigation, data curation, methodology, visualization, writing – review and editing

## 9. Declaration of generative AI and AI-assisted technologies in the writing process

During the preparation of this work the authors used ChatGPT (model 5.2)/Large language model in order to improve readability and language. After using this tool, the author reviewed and edited the content as needed and take full responsibility for the content of the published article.

## 14. Supplementary material

**Table S1:** Environmental gradient coverage by study period.

| Parameter | 1995–2009 | 1995–2009 | 2010–2025 | 2010–2025 |
| --- | --- | --- | --- | --- |
|  | min | max | min | max |
| Mean annual temperature (°C) | -2.56 | 7.64 | -1.16 | 8.2 |
| Atmospheric N deposition (mg m<sup>-2</sup> yr<sup>-1</sup>) | 63.2 | 1749.9 | 8.3 | 575.5 |
| Agriculture (% of catchment) | 0.0 | 44.9 | 0.007 | 44.9 |
| Forest (% of catchment) | 0.0 | 97.1 | 0.067 | 97.1 |
| Bog / peatland (% of catchment) | 0.005 | 23.0 | 0.032 | 23.0 |
| Lake color (proxy for DOM) | 1 | 111 | 2 | 242 |

**Table S2:** Spearman rank correlation matrix for predictor variables used to identify drivers of nitrogen-related limitation (study period 1995-2009). The numbers in the table are Spearman’s rank correlation coefficients (ρ, “rho”).

|  | Mean<br>annual<br>temperature | Atmospheric<br>N<br>deposition | Agriculture | Forest | Bog /<br>peatland | Lake color<br>(proxy for<br>DOM) |
| --- | --- | --- | --- | --- | --- | --- |
| Mean annual<br>temperature | 1.000 | 0.804 | 0.401 | 0.250 | -0.263 | 0.246 |
| Atmospheric<br>N deposition | 0.804 | 1.000 | 0.510 | 0.471 | -0.286 | 0.235 |
| Agriculture | 0.401 | 0.510 | 1.000 | 0.346 | 0.206 | 0.260 |
| Forest | 0.250 | 0.471 | 0.346 | 1.000 | 0.371 |  |
| Bog/ peatland | -0.263 | -0.286 | 0.206 | 0.301 | 1.000 | 0.234 |
| Lake color<br>(proxy for<br>DOM) | 0.246 | 0.235 | 0.260 | 0.371 | 0.234 | 1.000 |

**Table S3:**
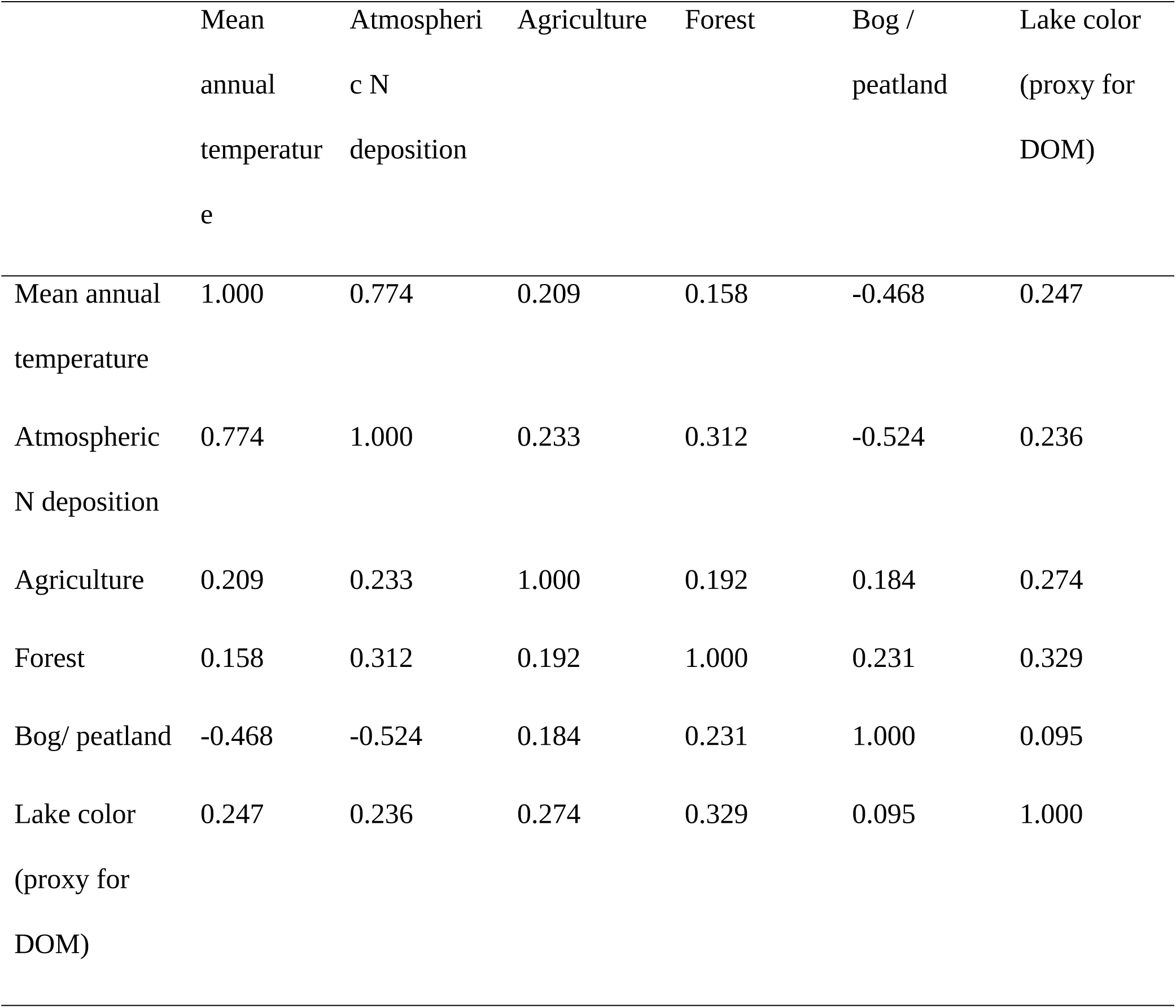
Spearman rank correlation matrix for predictor variables used to identify drivers of nitrogen-related limitation (study period 2010-2025). The numbers in the table are Spearman’s rank correlation coefficients (ρ, “rho”).

|  | Mean<br>annual<br>temperature | Atmospheric<br>N<br>deposition | Agriculture | Forest | Bog /<br>peatland | Lake color<br>(proxy for<br>DOM) |
| --- | --- | --- | --- | --- | --- | --- |
| Mean annual<br>temperature | 1.000 | 0.774 | 0.209 | 0.158 | -0.468 | 0.247 |
| Atmospheric<br>N deposition | 0.774 | 1.000 | 0.233 | 0.312 | -0.524 | 0.236 |
| Agriculture | 0.209 | 0.233 | 1.000 | 0.192 | 0.184 | 0.274 |
| Forest | 0.158 | 0.312 | 0.192 | 1.000 | 0.231 | 0.329 |
| Bog/ peatland | -0.468 | -0.524 | 0.184 | 0.231 | 1.000 | 0.095 |
| Lake color<br>(proxy for<br>DOM) | 0.247 | 0.236 | 0.274 | 0.329 | 0.095 | 1.000 |

**Table S4:**
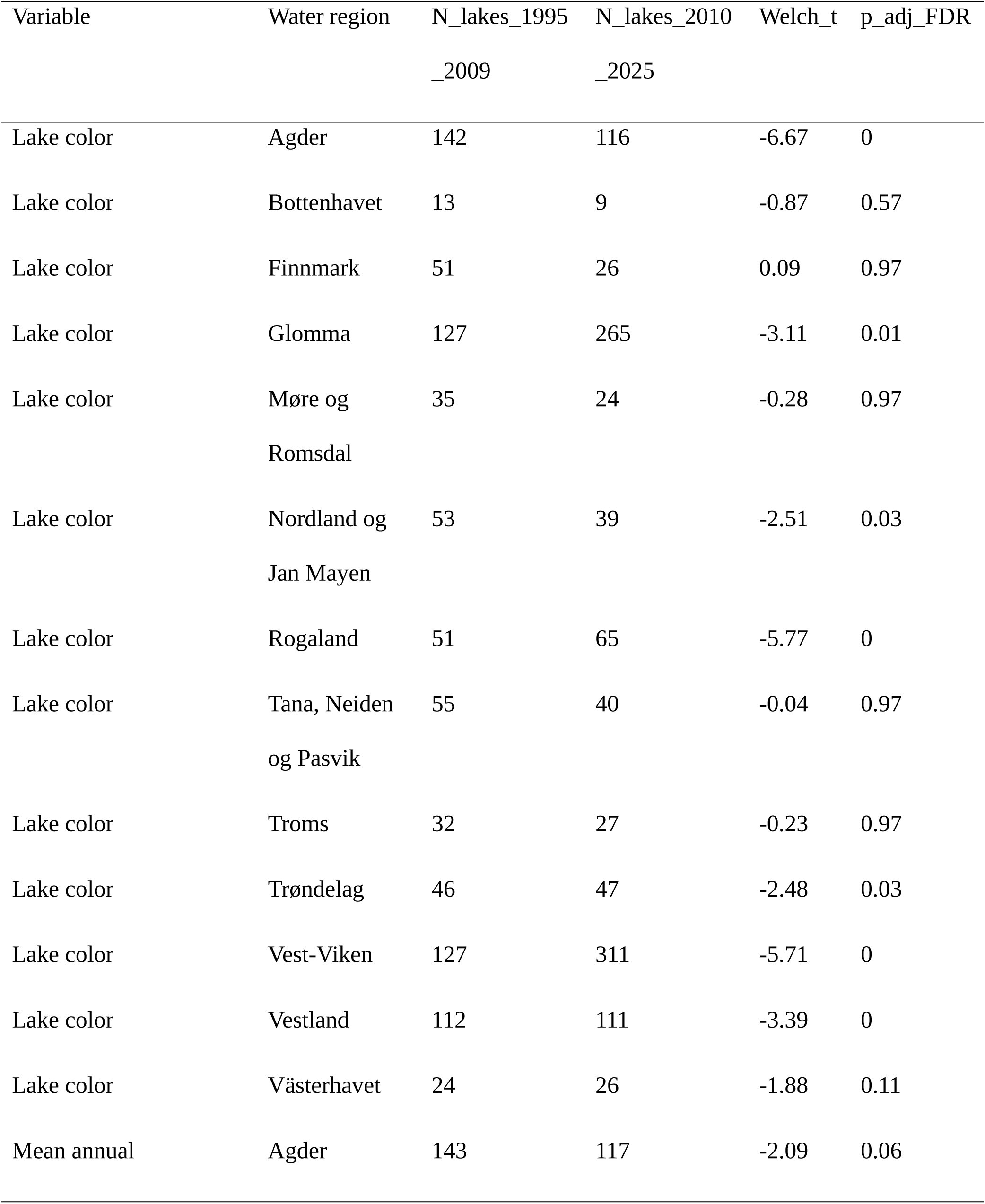
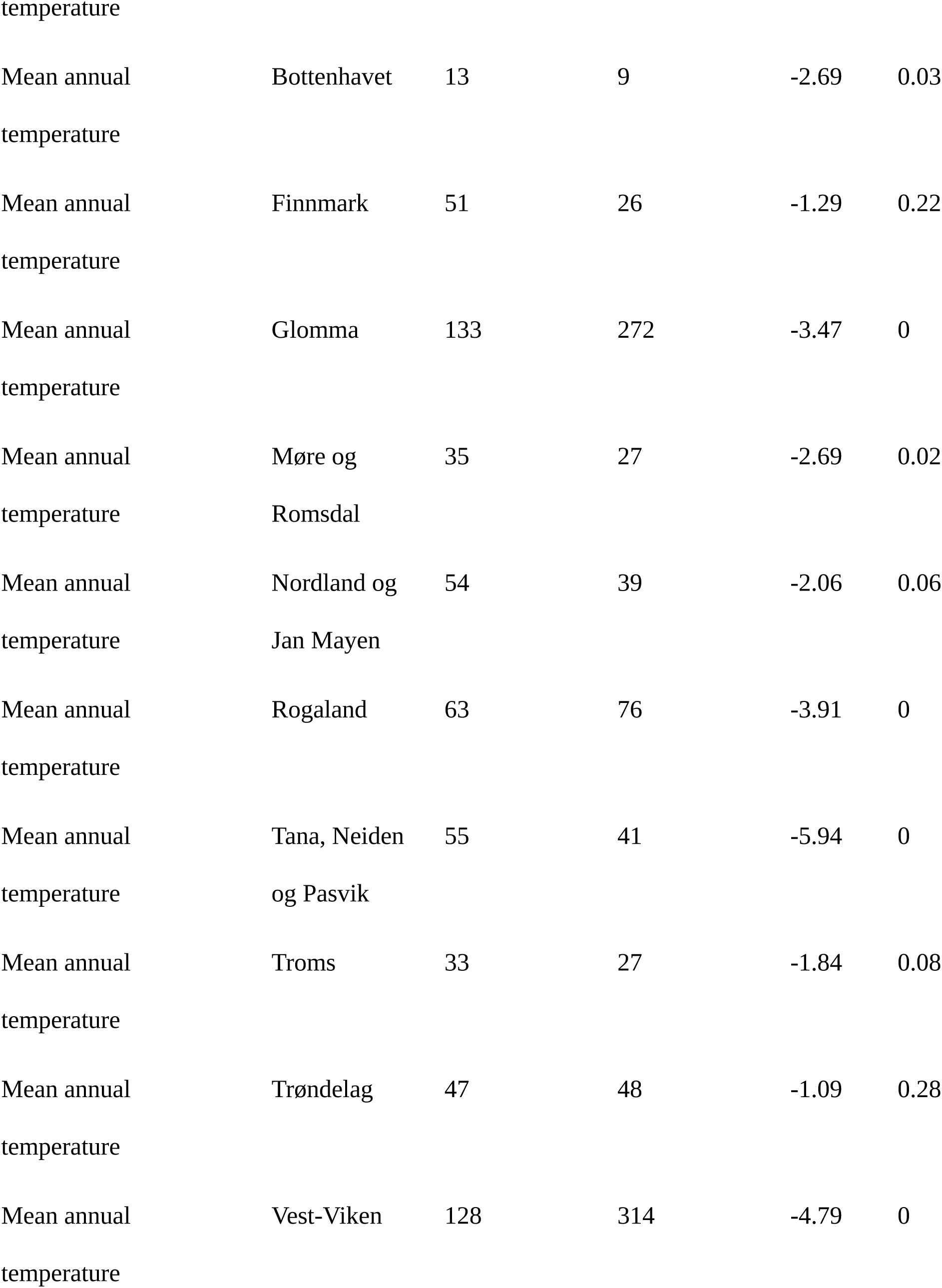
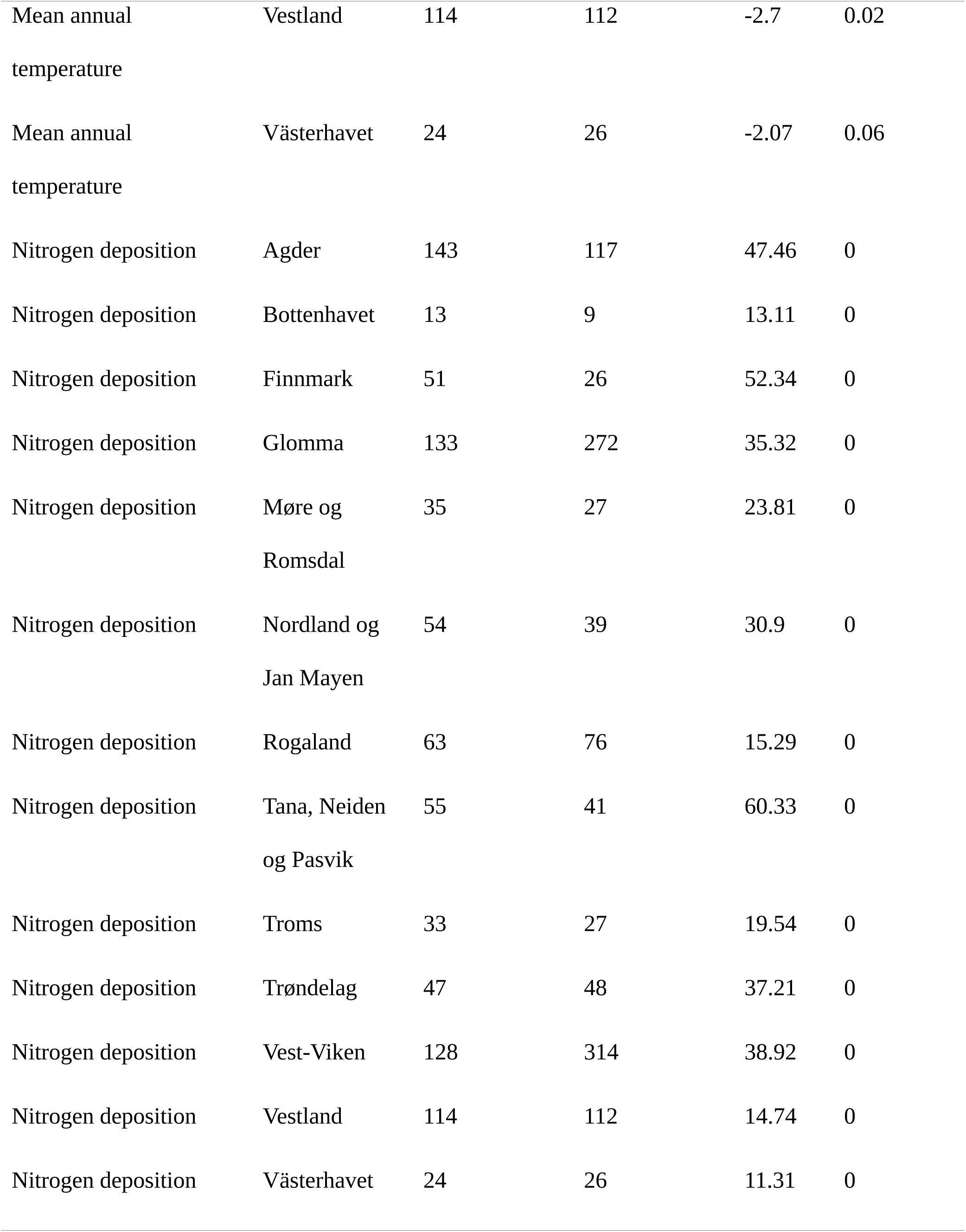
Results of statistical comparisons of predictor values between study periods.

**Table S5:**
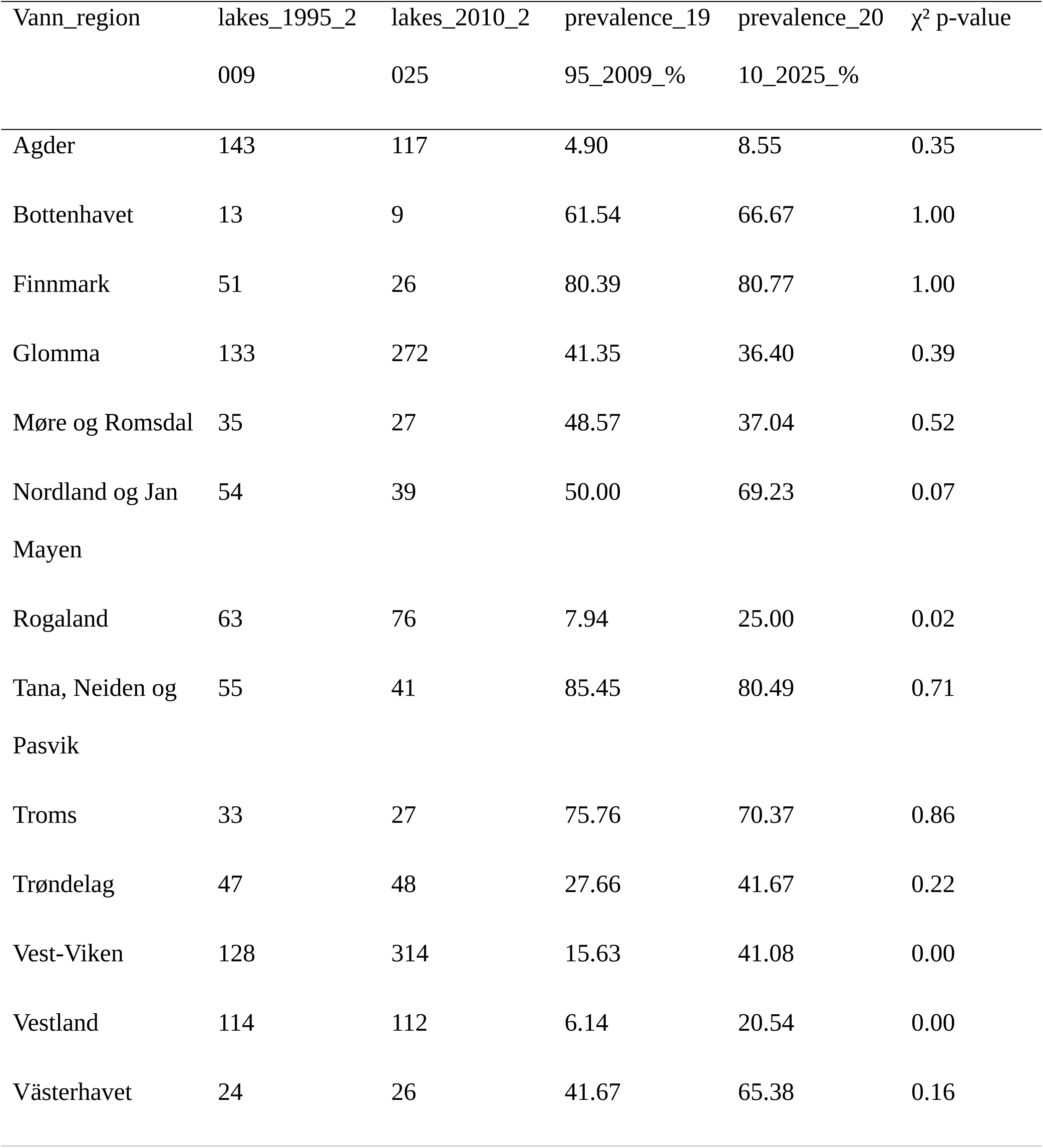
Results of statistical comparisons of prevalence of nitrogen-related limitation between study periods.

**Fig. S1:**
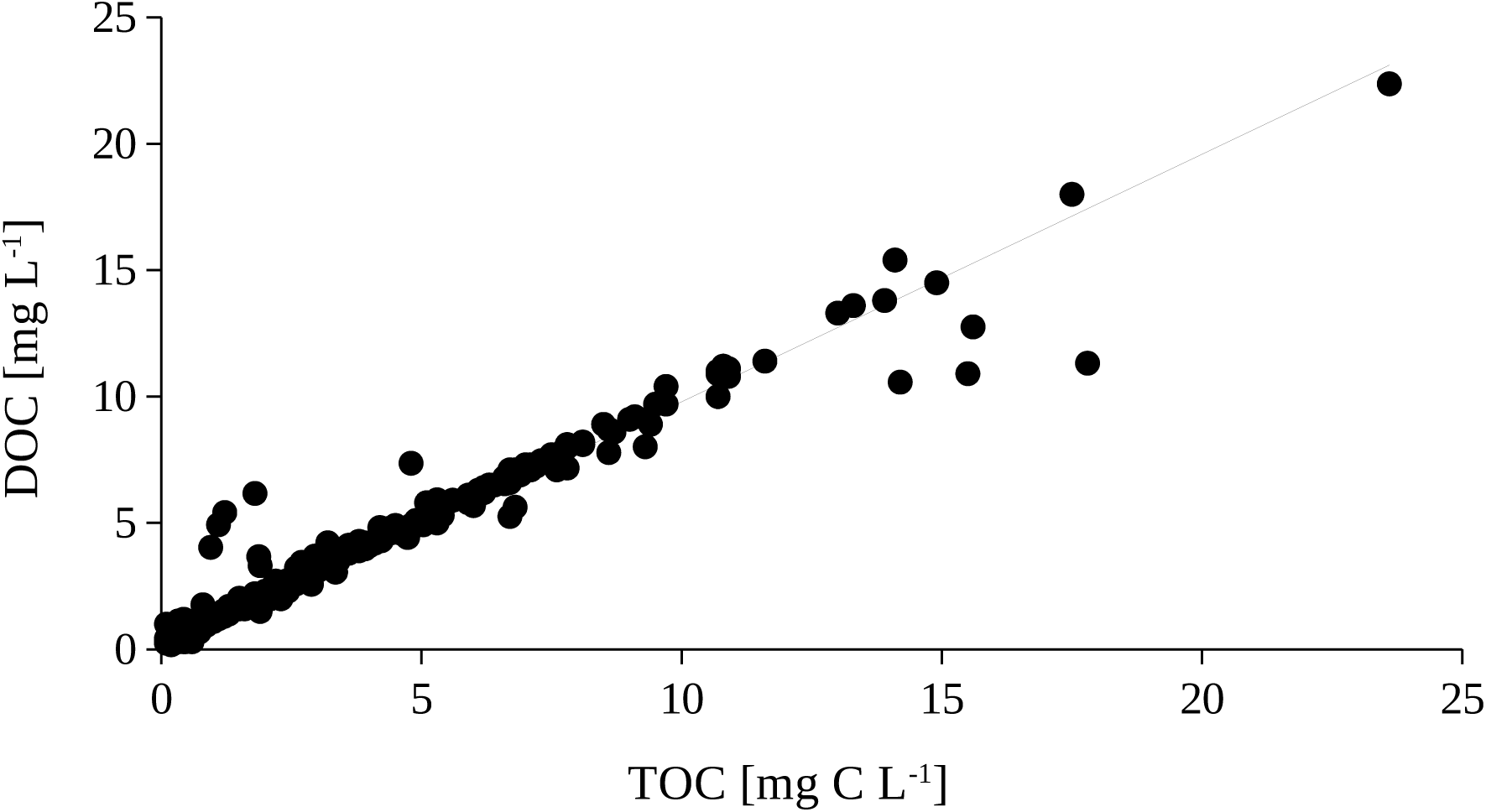
Linear regression between TOC and DOC across both study periods (R^2^=0.99, slope=1, p<0.0001).

**Fig. S2:**
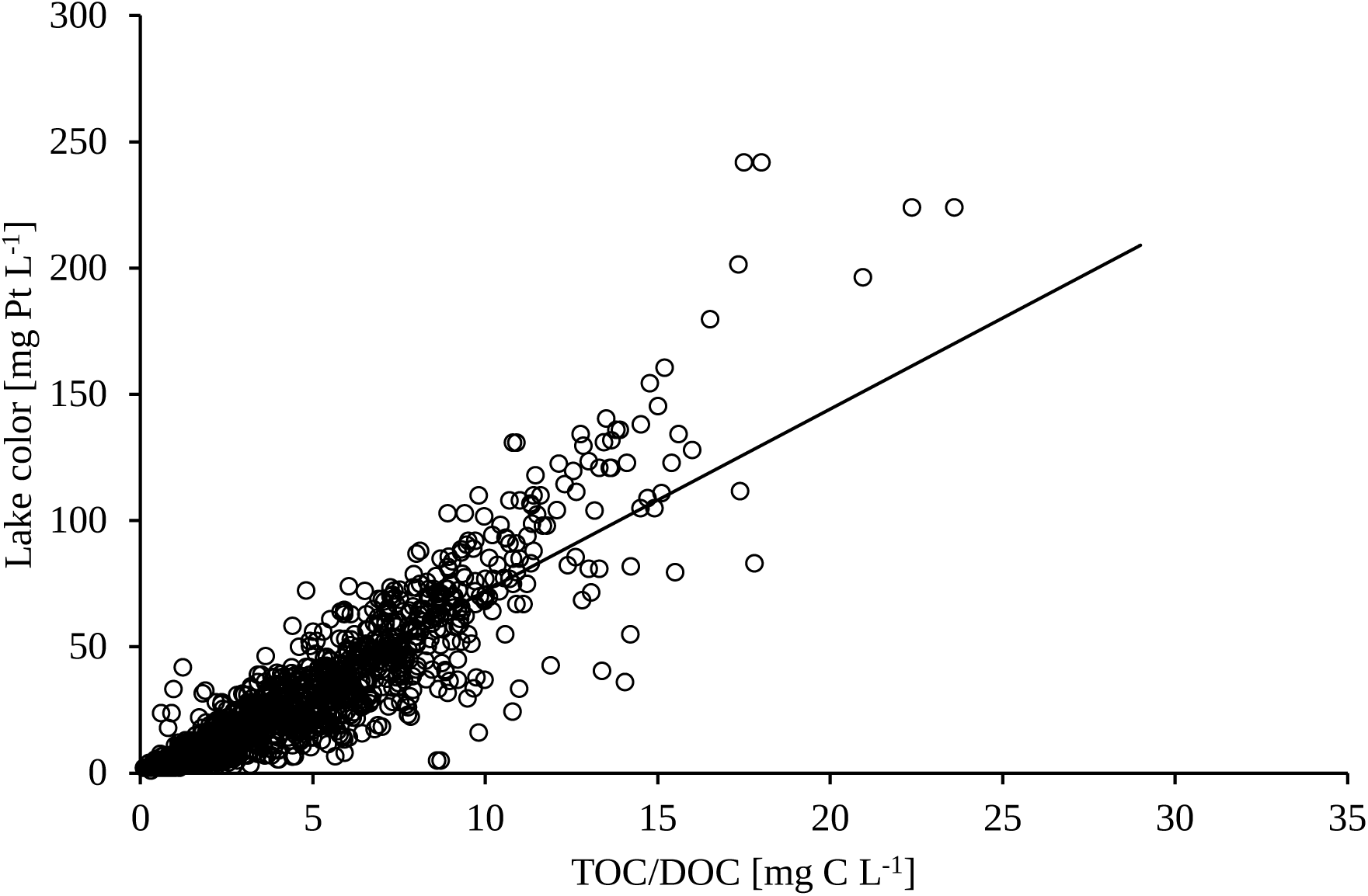
Linear regression between DOC/TOC and lake color across both study periods (R^2^=0.90, lake color=7.2 * TOC/DOC, p<0.0001). As shown in Fig. S1, DOC and TOC showed a linear relationship with a slope of 1 and were therefore both considered in this regression.

